# BET inhibitors as a therapeutic intervention in gastrointestinal gene signature-positive castration-resistant prostate cancer

**DOI:** 10.1101/2024.03.09.584256

**Authors:** Shipra Shukla, Dan Li, Holly Nguyen, Jennifer Conner, Gabriella Bayshtok, Woo Hyun Cho, Mohini Pachai, Nicholas Teri, Eric Campeau, Sarah Attwell, Patrick Trojer, Irina Ostrovnaya, Anuradha Gopalan, Eva Corey, Ping Chi, Yu Chen

## Abstract

A subgroup of castration-resistant prostate cancer (CRPC) aberrantly expresses a gastrointestinal (GI) transcriptome governed by two GI-lineage-restricted transcription factors, HNF1A and HNF4G. In this study, we found that expression of GI transcriptome in CRPC correlates with adverse clinical outcomes to androgen receptor signaling inhibitor treatment and shorter overall survival. Bromo- and extra-terminal domain inhibitors (BETi) downregulated HNF1A, HNF4G, and the GI transcriptome in multiple CRPC models, including cell lines, patient-derived organoids, and patient-derived xenografts, while AR and the androgen-dependent transcriptome were largely spared. Accordingly, BETi selectively inhibited growth of GI transcriptome-positive preclinical models of prostate cancer. Mechanistically, BETi inhibited BRD4 binding at enhancers globally, including both AR and HNF4G bound enhancers while gene expression was selectively perturbed. Restoration of HNF4G expression in the presence of BETi rescued target gene expression without rescuing BRD4 binding. This suggests that inhibition of master transcription factors expression underlies the selective transcriptional effects of BETi.

**SIGNIFICANCE:** GI transcriptome expression in CRPC is regulated by the HNF1A-HNF4G-BRD4 axis and correlates with worse clinical outcomes. Accordingly, BET inhibitors significantly reduce tumor cell growth in multiple GI-transcriptome-positive preclinical models of CRPC. Our studies point that expression of GI transcriptome could serve as a predictive biomarker to BETi therapy response.

## INTRODUCTION

Lineage plasticity is increasingly being appreciated as a mechanism to evade targeted therapy by cancer cells of multiple origins and lineages. Examples include prostate cancer and EGFR-mutant lung cancer where adenocarcinomas transdifferentiate into neuroendocrine cancers under select pressure of targeted therapy. In this process, cancer cells lose dependence on the initial tumor drivers, androgen receptor (AR) in prostate cancer, and EGFR and other oncogenic RTK in lung cancer (Sequist et al. 2011; Beltran et al. 2016; Quintanal-Villalonga et al. 2020). However, in contrast to a complete switch to neuroendocrine linage, a significant fraction of prostate adenocarcinoma also exists in a heterogeneous and plastic state where cancer cells acquire features of alternate cellular lineages and states such as stem cells, basal cells, mesenchymal cells, etc (Watson, Arora, and Sawyers 2015; Labrecque et al. 2019; Han et al. 2022; Tang et al. 2022). This poses a challenge in targeted therapy as 1) multiple dependencies exist in such tumors and 2) therapeutic targeting of the primary lineage may augment the process towards a complete lineage switch (Mu et al. 2017). Hence, combination therapies targeting more than one lineage/pathway may be more successful in such cases.

We have previously reported the activation of a gastrointestinal (GI) lineage transcriptome governed by aberrant expression of master regulators HNF1A and HNF4G in a significant fraction of castration-resistant prostate cancer (CRPC). HNF4G and HNF1A form a regulatory circuit where they influence each other’s expression. Exogenous expression of either HNF4G or HNF1A is sufficient to express the GI transcriptome in LNCaP cells that do not express either transcription factor. Expression of this aberrant GI transcriptome mediates resistance to enzalutamide (Shukla et al. 2017). In the present study, using two different metastatic CRPC (mCRPC) datasets, we show that increased GI transcriptome expression in patient tumors is associated with a shorter time on treatment with androgen receptor signaling inhibitors (ARSI) as well as a shorter overall survival. We hypothesized that inhibition of this transcriptome would provide therapeutic benefits in patients. Our studies revealed that inhibitors against Bromodomain and Extraterminal (BET) family member proteins efficiently inhibit GI transcriptome expression by directly targeting *HNF1A* and *HNF4G* transcription. Finally, we show the selective growth inhibitory effect elicited by BET inhibitors either alone or in combination with enzalutamide on GI transcriptome expressing preclinical CRPC models, including patient derived organoids and xenografts.

## RESULTS

### Aberrant expression of GI transcriptome in CRPC correlates with adverse clinical outcomes to ARSI treatment

The expression of GI transcriptome is governed by master regulators HNF1A and HNF4G and it is seen more prevalent in mCRPC compared to localized prostate cancer across multiple gene expression datasets (Shukla et al. 2017). Experimentally, exogenous expression of HNF4G in prostate cancer cells leads to expression of the GI transcriptome and resistance to AR pathway inhibition. These data suggest a causal relationship between the expression of GI transcriptome and resistance to AR-targeted therapy (Shukla et al. 2017).

Here, we sought to quantify the level of GI transcriptome expression and correlate it with clinical outcomes. We derived an *HNF signature* comprised of HNF1A, HNF4G, and their nine strong direct downstream targets and an *HNF score* derived from the summed z-scores of their gene expression. Correlation analysis performed on two clinical gene expression datasets showed that the HNF score is significantly correlated with HNF1A and HNF4G expression (**Figure S1A-B**). The HNF score also strongly correlated with the broader prostate cancer-gastrointestinal (PCa_GI) signature sum Z-score (**Figures S1A-B**). The PCa_GI signature is previously defined and derived from correlation with SPINK1 in primary prostate cancer (Shukla et al. 2017). We applied HNF score to analyze two RNA-Seq datasets of CRPC tumors from patients treated with ARSIs. The clinical trial Genetic and Molecular Mechanisms in Assessing Response in Patients with Prostate Cancer Receiving Enzalutamide Therapy (NCT02099864) prospectively enrolled 36 taxane and abiraterone naïve mCRPC patients to treatment with enzalutamide (Alumkal et al. 2020). Response was defined as a 50% decline in PSA after 12 weeks of treatment. Among the 25 patients with pre-treatment RNA-Seq data, we found that 20% of tumors (n=5) had a higher HNF score (*Z*>12) than the rest. We used this cutoff to define HNF score_High tumors (**Figure 1A**). Notably, four of five patients with HNF score_High tumors but only one of twenty patients with HNF score_Low tumors did not respond to enzalutamide treatment (Fisher’s exact test P=0.012) (**Figure 1B**). Alternatively, 4/7 non-responders showed a high HNF score as compared to 1/18 of responders (**Figure S1C**). Global transcriptome analysis showed a significant upregulation of many GI lineage genes such as HNF1A, MUC13, UGT2B4, MIA2, and NR1H4 in enzalutamide non-responders compared to responders (**Figure S1D)**. We performed Gene Set Enrichment Analysis (GSEA) comparing non-responders and responders using the Molecular Signatures Database (MSigDB) comprising >20,000 gene sets and our custom gene sets. We found our previously defined PCa_GI_signature gene set as well as a gene set comprised of HNF1A targets to be significantly enriched in enzalutamide non-responders (**Figure 1C**). We next analyzed the RNA-Seq data of mCRPC patients from SU2C International Dream Team and calculated the HNF score for patients for whom the overall survival and time on ARSI treatment was available (Robinson et al. 2015; Abida et al. 2019). We analyzed ARSI naïve patients going onto ARSI therapy (n=50).

**Figure 1.**
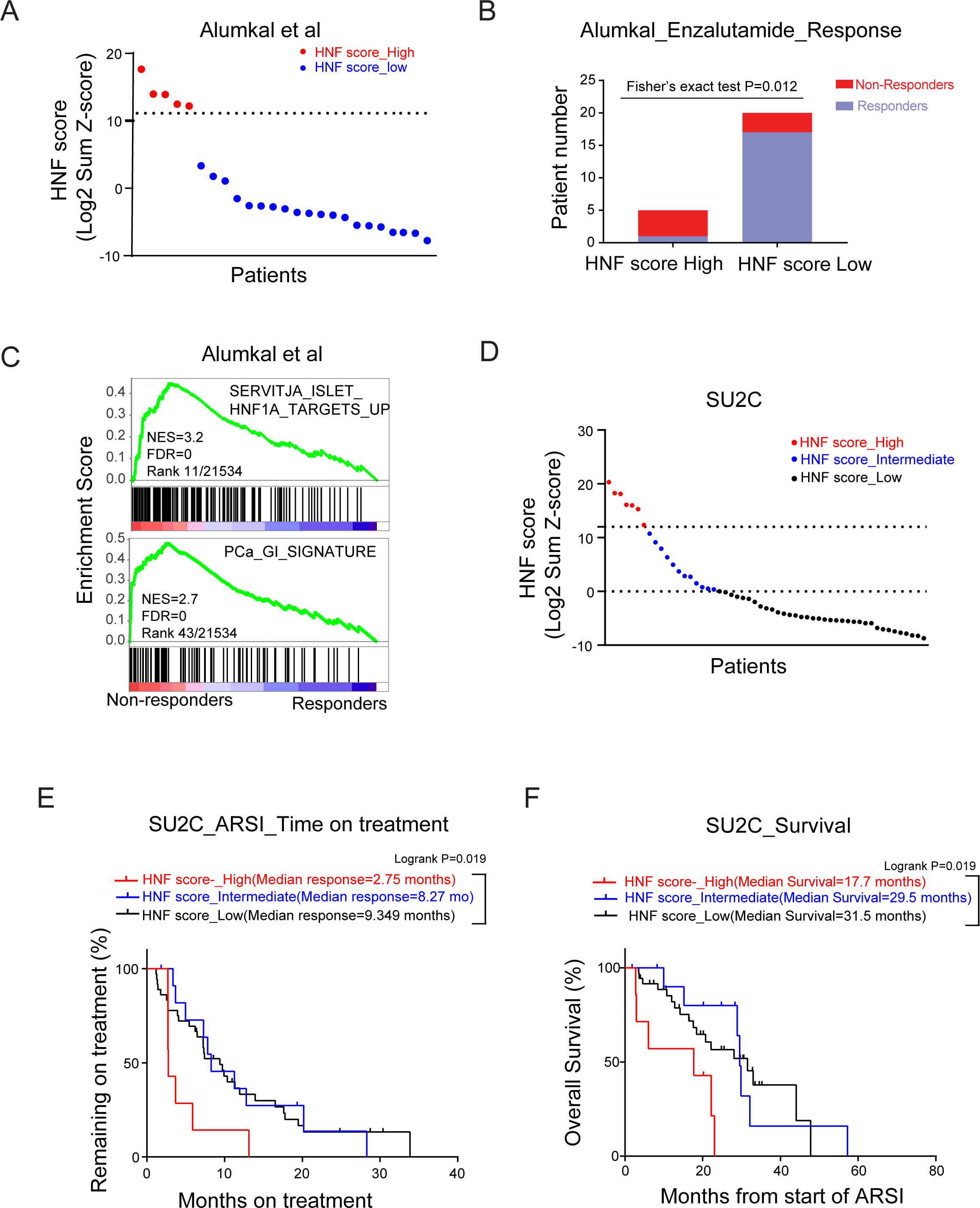
A high HNF score in CRPC correlates with adverse clinical outcomes. (A) Patient stratification based on HNF scores in the Alumkal dataset. Each dot represents one patient. HNF score was calculated as the log2 sum z-score of mRNA expression of 11 genes. A sum z-score of >=12 was annotated as a high HNF score and <12 as a low HNF score. See methods for details. (B) Enzalutamide response of patient tumors with high and low HNF scores. Statistical significance is determined using the Fisher’s exact test. (C) GSEA plots of HNF1A target genes (top) and PCa_GI Gene signature (bottom) in enzalutamide non-responders compared to responders. NES: Normalized enrichment score. FDR: False discovery rate. (D) Patient stratification based on HNF score expression in SU2C dataset. Each dot represents one patient. Tumors with a sum z-score of >=12 were annotated as expressing high HNF score; <=0 as low HNF score and a value between 0-12 as intermediate HNF_score. See methods for details. (E) Kaplan–Meier curve comparing ARSI outcome measures between the three groups stratified by HNF scores. P values were determined using the log-rank test. (F) Kaplan–Meier curve comparing overall survival outcome between the three groups stratified by HNF scores. P values were determined using the log-rank test.

Ranking of patients based on the HNF scores annotated them into three categories: patients with a sum z-score value of >12 as in Alumkal dataset were categorized as HNF score_High while patients with a sum z-score of zero or less were categorized as HNF score_Low. The remaining patients were categorized as HNF score_Intermediate (**Figure 1D**). Kaplan Meier analysis revealed that the patients categorized as HNF score_High had the shortest median time on ARSI (**Figure 1E**). To investigate whether the poor response to ARSI would translate to shorter overall survival of these patients, we performed a Kaplan-Meier survival analysis and found that HNF score_High patients had a significantly shorter overall survival as compared to the other two cohorts (**Figure 1F**). These data suggest that increased expression of the GI transcriptome in patients is associated with worse clinical outcomes in CRPC patients.

### BET inhibition downregulates GI transcriptome in CRPC

Previously, we showed that HNF4G is required for maintaining open chromatin regions and active transcription-associated epigenetic modifications such as H3K4me1 and H3K27ac at its target genes. Members of the BET family proteins, BRD2, BRD3, and BRD4 are epigenetic readers. They bind to acetylated histones through their bromodomains and facilitate the assembly of active transcriptional complexes. In the absence of selective inhibitors against HNF1A and HNF4G, we explored the impact of targeting BET proteins on GI transcriptome expression. We used two BET inhibitors, ABBV-075 (mivebresib) and JQ1, in experiments performed on 22Rv1 cells that express the GI transcriptome. Treatment with either inhibitor for 4 hours led to a dose-dependent decrease in transcripts of HNF1A and HNF4G (**Figure 2A, B**), while the AR transcript was only modestly inhibited at high concentrations (**Figure S2A**). Immunoblot analysis at 24 hours post-treatment, showed reduced protein levels of HNF1A and HNF4G, as well as their downstream targets AKR1C3 and UGT2B15, indicating that BET proteins regulate the transcription of these genes (**Figure 2C**). To analyze the effect of BET inhibition on global gene expression, we performed RNA-Seq of 22Rv1 cells treated with 25 nM ABBV-075 for 24 hours. The ABBV-075 treatment led to a downregulation of the HNF signature as well as the broader PCa_GI signature (**Figures 2D and S2B)**. However, the effect of ABBV-075 on the AR transcriptome varied, as assessed using two different AR signatures (Bluemn et al. 2017; Hieronymus et al. 2006) (**Figure 2D**). We next performed GSEA on RNA-Seq data obtained from DMSO and ABBV-075 treated cells. The topmost downregulated gene sets in ABBV-075 treated cells included the PCa_GI signature and gene sets regulated by HNF4G and HNF1A (**Figure 2E**).

**Figure 2.**
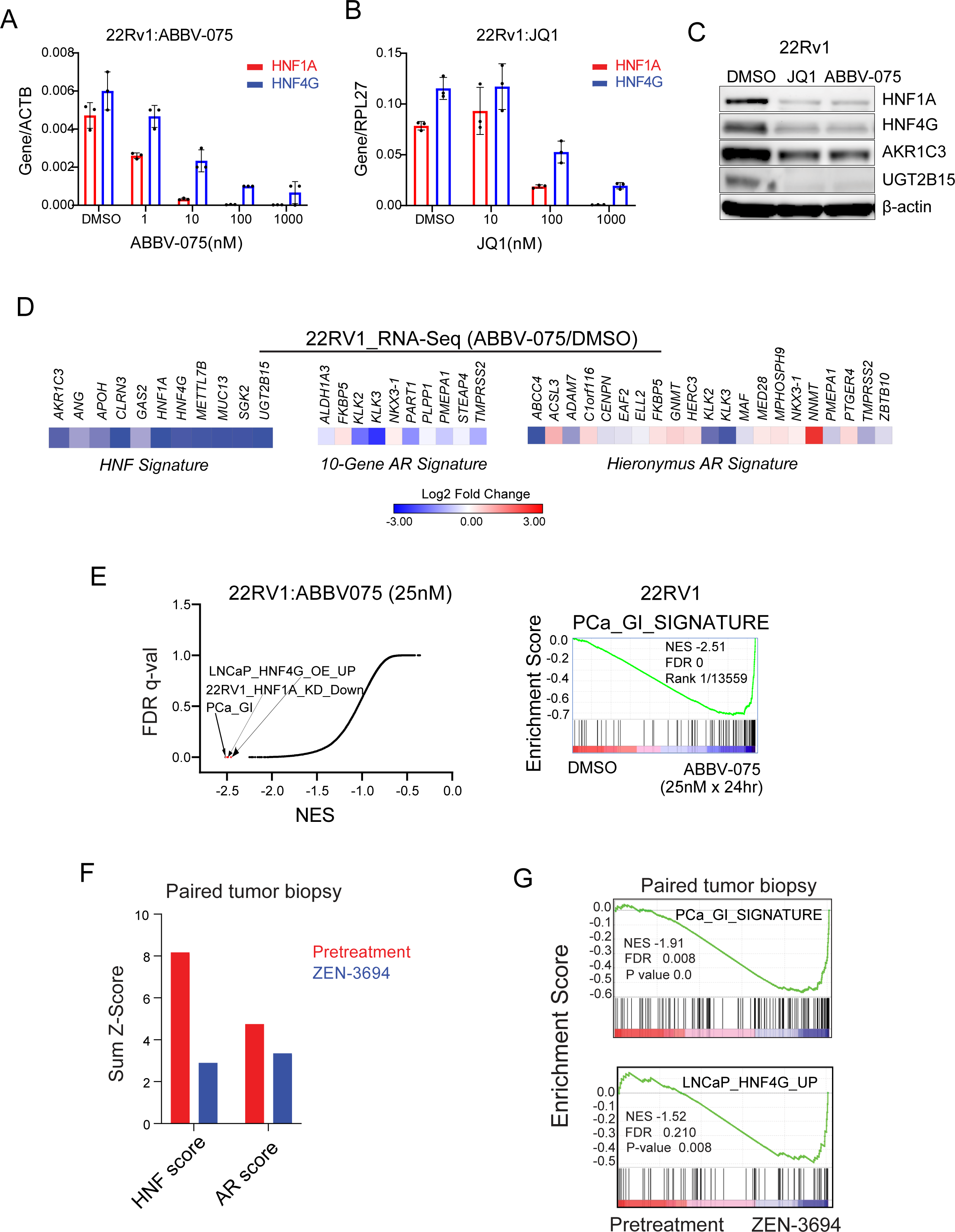
BET inhibitors downregulate the expression of HNF4G and HNF1A and their transcriptional signature. (A) qRT-PCR showing expression of HNF1A after 4 hours of treatment with ABBV-075 and JQ1 at indicated doses. (B) qRT-PCR showing expression of HNF4G after 4 hours of treatment with ABBV-075 and JQ1 at indicated doses. (C) Immunoblots of 22Rv1 cells treated with JQ (1 µM), ABBV-075 (50 nM), and DMSO control for 24 hours against the indicated proteins. (D) Heatmap of RNA-Seq expression of HNF signature genes in 22Rv1 cells after treatment with 25 nM ABBV-075 for 24 hours (left). The middle and right heatmaps show the modulation of AR target genes with ABBV-075 treatment using two different AR gene signatures. Data is plotted as the log2 difference in gene expression between ABBV-075 and DMSO treated cells. (E) Global representation of GSEA analysis of RNA-Seq gene expression data set of 22RV1 cells treated with 25 nM ABBV-075 for 24 hours. X-axis shows the normalized enrichment score, and y-axis is the FDR q-value. The PCa_GI and the HNF1A, and HNF4G regulated gene sets are indicated in red. GSEA plot of broader PCa_GI gene signature is shown in the right. NES: Normalized enrichment score. FDR: False discovery rate. (F) Modulation of HNF and AR scores by BETi ZEN-3694 in paired tumor biopsies of patient 101047. (G) GSEA plots of PCa_GI Gene signature (top) and HNF4G target genes (bottom) in ZEN-3694 treated tumors compared to pretreated tumor. NES: Normalized enrichment score. FDR: False discovery rate.

In addition, we analyzed publicly available data on BET inhibition (JQ1 and ABBV075) in 22Rv1 cells (Cai et al. 2018; Faivre et al. 2020; Welti et al. 2021). GSEA using RNA-Seq data from DMSO, JQ1, and ABBV-075 treated cells showed that, for both JQ1 and ABBV-075, gene sets regulated by HNF1A and HNF4G, and the PCa_GI signature, were among the topmost significantly downregulated gene sets. Consistent with our observations, BETi treatment caused downregulation of HNF1A and HNF4G but not AR (**Figure S2C-E**).

BET inhibitors, including ZEN-3694 and NUV-868 are being evaluated in clinical trials in various cancer types, including prostate cancer (NCT02705469, NCT04471974, NCT04986423, NCT02711956, NCT05252390). We took advantage of a recently completed Phase 1b/2a clinical trial of the BET inhibitor, ZEN-3694 on a cohort of mCRPC (Aggarwal et al. 2020). Gene expression data from pre- and post-ZEN-3694 treatment was available for four patients. Among them, pre-treatment biopsy from patient 101047 exhibited a high HNF score. RNA-Seq analysis performed on the paired biopsies of patient 101047 tumors revealed downregulation of HNF score post-ZEN-3694 treatment compared to the pretreatment biopsy. While the AR score was only modestly downregulated upon BET inhibition (**Figure 2F**). GSEA showed the downregulation of the PCa_GI_signature and HNF4G target gene sets by ZEN-3694 in the post-treatment biopsy (**Figure 2G**). ZEN-3694 treatment caused downregulation of HNF1A and HNF4G but not AR in the post-treatment biopsy (**Figure S2F**).

### Inhibition of HNF4G transcription principally accounts for BETi-mediated inhibition of GI transcriptome

We asked whether the preferential inhibition of GI transcriptome over AR-regulated transcriptome by BETi treatment is due to downregulation of master transcription factors HNF4G and HNF1A but not of AR. To explore this possibility, we generated 22Rv1 derivatives that exogenously express HNF4G (HNF4G OE) or GFP (GFP OE) from the Murine Stem Cell Virus (MSCV) promoter that is not repressed with ABBV-075 (**Figure S3A-B)**. We then treated GFP OE and HNF4G OE cells with ABBV-075 (25 nM) or DMSO for 24 hours and performed RNA-Seq analysis. We compared the effect of ABBV-075 treatment on the expression of 31 direct targets of HNF4G between the GFP or HNF4G expressing cells using DMSO treatment as a control. We observed that restoring the expression of HNF4G can largely reverse the ABBV-075-mediated downregulation of its direct targets (**Figure 3A**). Examination of individual genes shows partial (*HNF1A*) to almost complete (*CCN2*, *CLRN3*, *VIL1*) rescue of transcriptional inhibition (**Figure 3B**).

**Figure 3.**
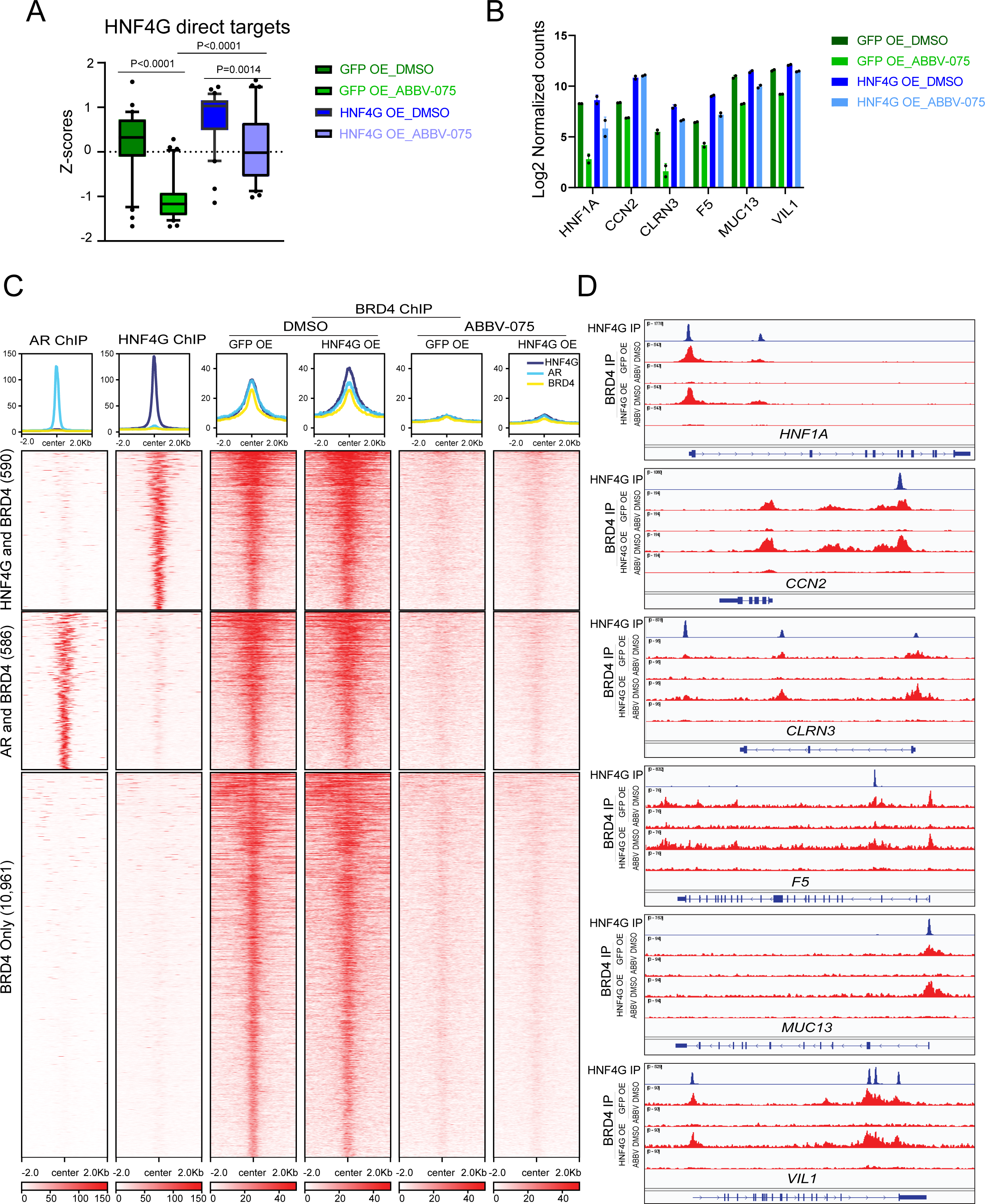
Inhibition of HNF4G transcription accounts for BETi mediated inhibition of GI transcriptome. (A) Box plot of changes in Z-score expression values of HNF4G direct targets by ABBV-075 treatment in 22Rv1 cells exogenously expressing GFP or HNF4G compared with DMSO control. Box plots show median, quartiles, min, and max, with 10-90 percentile dots plotted. Statistical analysis was performed using a two-tailed unpaired t-test. (B) RNA-Seq gene expression values of selected HNF4G target genes after exogenous expression of HNF4G and GFP in 22Rv1cells treated with DMSO or ABBV-075 (25 nM). Data are presented as mean ± SD. (C) Histograms (top) show the average normalized tag counts of AR and HNF4G in parental 22Rv1 cells and that of BRD4 in GFP or HNF4G expressing 22Rv1 cells treated with ABBV-075 or DMSO at top 1,000 HNF4G, 1,000 AR binding sites and BRD4 only enhancer binding sites. Heatmap shows the tag densities of HNF4G, AR, and that of BRD4 at HNF4G (top) or AR (middle) binding sites. Bottom panel show the tag densities of BRD4 at 10,961 BRD4 only sites in GFP or HNF4G expressing 22Rv1 cells treated with ABBV-075 or DMSO. (D) ChIP-seq profiles of HNF4G in parental 22Rv1 cells and BRD4 (DMSO treatment), and BRD4 (ABBV-075 treatment) in GFP or HNF4G expressing 22Rv1 cells at selected HNF4G target genes loci; *HNF1A*, *CCN2*, *CLRN3*, *F5*, *MUC13*, and *VIL1* in top to bottom order.

Since BRD4 is the most extensively characterized member of BET family proteins, we next examined the requirement of BRD4 at the loci of these transcriptionally rescued genes. We performed BRD4 ChIP-Seq in GFP OE and HNF4G OE cells treated with ABBV-075 (25 nM for 4 hours) or DMSO. We examined BRD4 binding at BRD4 peaks that overlapped with previously defined top 1,000 HNF4G peaks (n=590), top 1,000 AR peaks (n=586), and non-overlapping BRD4 peaks (n=10,961). Exogenous expression of HNF4G led to a modest increase of BRD4 binding at HNF4G binding sites but not at AR binding sites or non-overlapping sites. ABBV-075 treatment broadly displaced BRD4 from chromatin at all BRD4 peaks and exogenous HNF4G expression did not rescue BRD4 binding (**Figure 3C**). Examination of ChIP-Seq tracks of selected genes shown in figure 3B reveals a similar level of BRD4 displacement with ABBV-075 treatment in between GFP OE and HNF4G OE cells despite their continued transcription in HNF4G OE cells (**Figure 3D**). These data suggest that BRD4 is an accessory factor rather than the primary factor in controlling gene expression regulation. At its target genes, restoration of HNF4G expression mitigates the transcriptional effects of BRD4 displacement. More broadly, these data suggest that the downregulation of master transcription factors underlies the selectivity of BETi transcriptional inhibition despite the broad displacement of BET proteins from chromatin (Cai et al. 2018).

### GI-transcriptome-positive prostate cancer models exhibit increased sensitivity to BET inhibitors

To examine the effect of BET inhibition on the growth of GI transcriptome-positive prostate cancer, we treated ten prostate cancer organoids derived from mCRPC patients with ABBV-075 (Gao et al. 2014; Tang et al. 2022). Notably, we observed that organoids with a high HNF score; MSK-PCa17, MSK-PCa13, and MSK-PCa10, exhibited the lowest half-maximal inhibitory concentration (IC50) to ABBV-075 treatment, suggesting high sensitivity (**Figures 4A and S4A**). MSK-PCa17 cells had the lowest IC50 (IC50<2 nM) among all the organoids.

**Figure 4.**
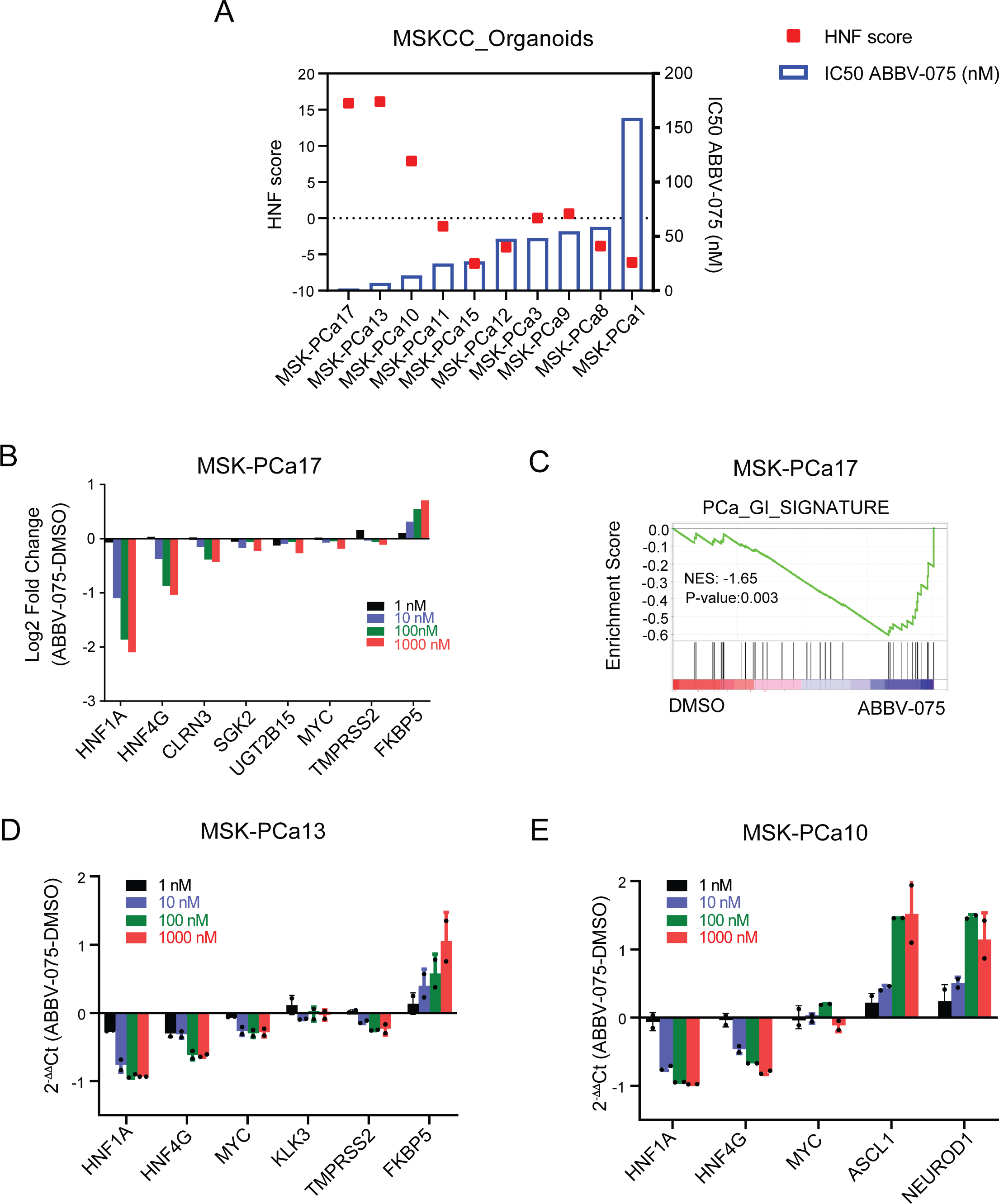
Patient-derived organoids with high HNF scores show increased sensitivity to BETi-mediated growth inhibition. (A) IC50 of ABBV-075 in a panel of patient-derived tumor biopsies grown as organoids. The left Y-axis plots the HNF scores of each organoid and the right Y-axis shows the IC50 values. (B) RNA-Seq gene expression changes of selected genes at different doses of ABBV-075 treatment of MSK-PCa17 cells compared to DMSO control. Data is presented as log2 fold difference in expression (ABBV-075 vs DMSO). (C) GSEA analysis indicating the negative enrichment of PCa_GI gene signature gene set in MSK-PCa17 cells treated with ABBV-075 (10 nM) compared to DMSO control. NES: Normalized enrichment score. (D) qRT-PCR showing expression of selected genes after 4 hours of treatment with ABBV-075 at indicated doses in MSK-PCa13 cells. (E) qRT-PCR showing expression of selected genes after 4 hours of treatment with ABBV-075 at indicated doses in MSK-PCa10 cells.

To identify important genes/pathways perturbed by ABBV-075, we performed RNA-Seq on MSK-PCa17 cells treated with ABBV-075 at four different concentrations (1nM, 10 nM, 100 nM, and 1000 nM) for four hours. Our results showed that *HNF1A* and *HNF4G* were downregulated in a dose-dependent manner, along with other signature GI transcriptome genes such as CLRN3, SGK2, and UGT2B15 (**Figure 4B**). GSEA showed that the PCa_GI_signature was downregulated upon ABBV-075 treatment in MSK-PCa17 (**Figure 4C**). In MSK-PCa13 cells, the second most sensitive line, qRT-PCR analysis demonstrated a dose-dependent decrease in HNF1A and HNF4G transcript levels with ABBV-075 treatment (**Figure 4D**). MSK-PCa10, a neuroendocrine prostate cancer (NEPC) organoid with a high HNF score, showed sensitivity to BET inhibition. qRT-PCR analysis revealed no decrease in important NEPC lineage genes such as ASCL1 and NEUROD. However, HNF1A transcription was strongly suppressed by ABBV-075 in these cells (**Figure 4E**). These data suggest that organoids with a high HNF score are sensitive to growth inhibition by ABBV-075, emphasizing the potential relevance of the GI transcriptome expression to BET inhibition response. The observed downregulation of key genes and pathways associated with the GI transcriptome supports the potential therapeutic efficacy of BET inhibition in this context.

Next, we assessed response to BETi in vivo using a panel of twelve LuCaP patient-derived xenografts (PDXs). The LuCaP PDXs are well annotated and represent the varied clinical spectrum of CRPC (Lam et al. 2017; Nguyen et al. 2017). We chose pelabresib (CPI-0610) for in vivo studies because it has favorable pharmacokinetics and pharmacodynamics properties, and it is in late-stage clinical development (Albrecht et al. 2016). We treated each PDX with pelabresib or vehicle for four weeks. Fold changes in tumor volume were determined by comparing the pelabresib-treated group to the vehicle-treated group after the four-week treatment period. HNF scores for each PDX were calculated using baseline RNA-Seq data. We observed that PDXs with higher HNF scores were more sensitive to the growth inhibitory effects of pelabresib (**Figure 5A**). The data suggest that the GI transcriptional activity, as assessed by HNF score, may serve as a predictive marker for the responsiveness of prostate cancer PDXs to pelabresib treatment. The observed correlation highlights the potential clinical relevance of the GI transcriptome in guiding BET inhibitor therapy.

**Figure 5.**
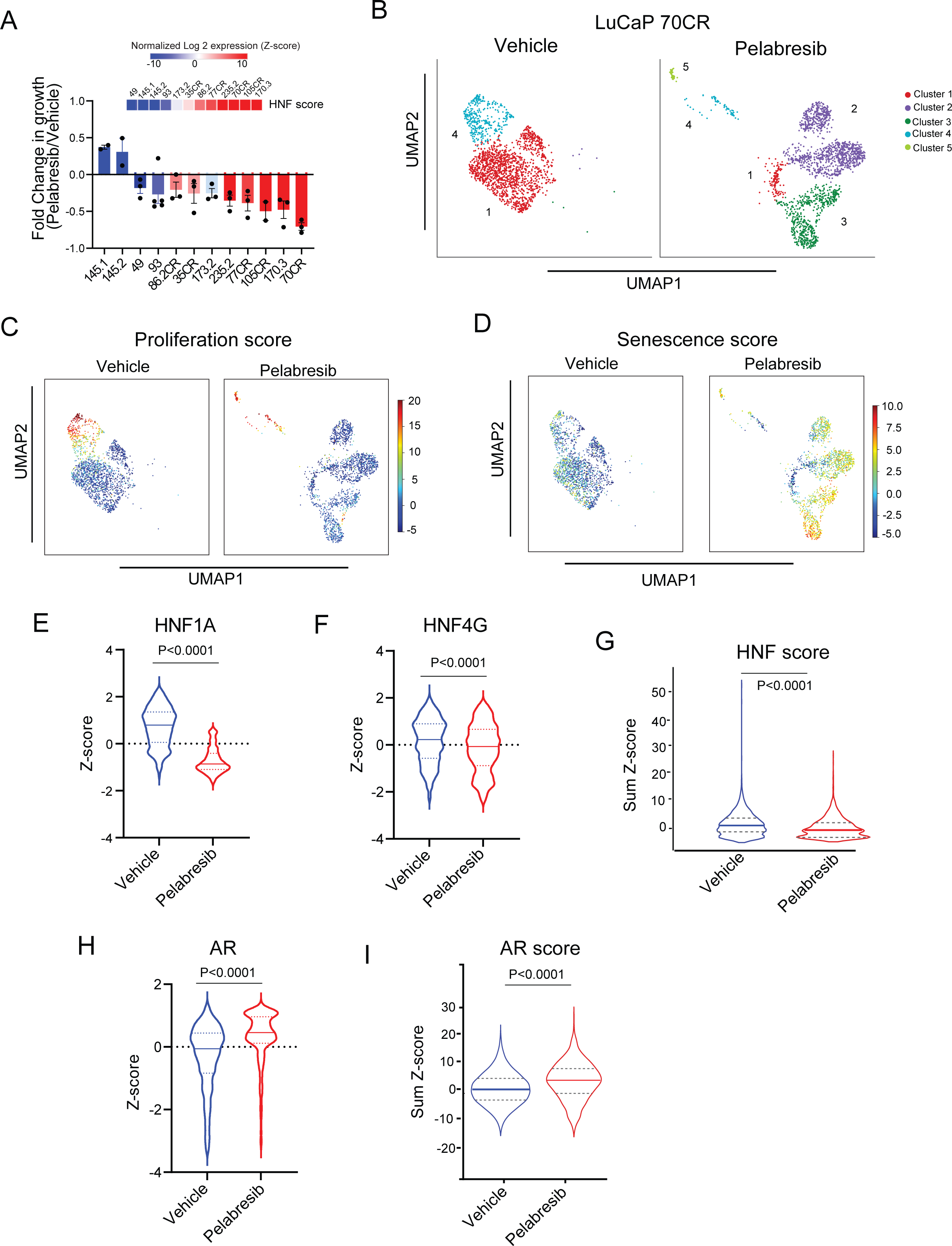
CRPC PDXs expressing high HNF score are sensitive to BET inhibition. (A) Treatment response of LuCaP PDXs when treated with pelabresib (30 mg/kg) or vehicle (1% carboxymethyl cellulose) twice a day. Treatment was started when tumors reached a volume of approximately 100 mm^3^. Data is plotted as the fold change in tumor volume between pelabresib and vehicle treated tumors after 4 weeks of treatment. Mean ± SEM. Two-tailed unpaired t*-*test, n=3. HNF score of PDXs calculated using the 11 gene signature is shown on top of the graph. (B) UMAPs of single cells isolated from vehicle or pelabresib treated LuCaP 70CR tumors. (C) UMAPs depicting the proliferation scores of single cells isolated from vehicle or pelabresib treated LuCaP 70CR tumors. (D) UMAPs depicting senescence scores of single cells isolated from vehicle or pelabresib treated LuCaP 70CR tumors. (E) Violin plot of HNF1A expression in single cells obtained from vehicle and pelabresib treated tumors. The median is shown by a solid line while the first and third quartiles are shown by dashed lines. P value is obtained from unpaired t-test. (F) Violin plot of HNF4G expression in single cells obtained from vehicle and pelabresib treated tumors. The median is shown by a solid line while the first and third quartiles are shown by dashed lines. P value is obtained from unpaired t-test. (G) Violin plot depicting HNF score in single cells obtained from vehicle and pelabresib treated tumors. The median is shown by a solid line while the first and third quartiles are shown by dashed lines. P value is obtained from unpaired t-test. (H) Violin plot depicting AR expression in single cells obtained from vehicle and pelabresib treated tumors. The median is shown by a solid line while the first and third quartiles are shown by dashed lines. P value is obtained from unpaired t-test. (I) Violin plot depicting AR score in single cells obtained from vehicle and pelabresib treated tumors. The median is shown by a solid line while the first and third quartiles are shown by dashed lines. P value is obtained from unpaired t-test.

To gain a comprehensive understanding of the molecular changes induced by BET inhibition, we performed single-cell RNA sequencing (scRNA-Seq) on LuCaP 70CR tumors treated with pelabresib for six days using vehicle as control. Tumors were dissociated into single-cell suspension and live cells were obtained using fluorescence-activated cell sorting. We discarded cells with mouse reads and analyzed single transcriptomes from approximately 3650 single human cells in vehicle and pelabresib-treated mice (n=2) after quality control and filtering. Dimension reduction using Uniform Manifold Approximation and Projection (UMAP) and Leiden clustering grouped tumor cells into five clusters (**Figure 5B**). In vehicle-treated mice, the majority of tumor cells grouped into cluster 1 that have characteristics of prostate adenocarcinoma including luminal markers KRT8, KRT18, FOLH1; prostate transcription factors AR, NKX3-1, FOXA1, HOXB13; and GI transcription factors HNF1A, HNF4G and downstream target like MUC13 (**Figure S5A**). In addition, a fraction of cells grouped into cluster 4, which maintained prostate lineage markers and are additionally characterized by expression of proliferation genes, suggesting this is the proliferative cluster (**Figures 5C and S5B-C**). Treatment with pelabresib resulted in a decrease in clusters 1 and 4 cell population and an increase/emergence of clusters 2, 3, and 5 (**Figure 5B**). These 3 clusters all expressed senescence-related genes and exhibited a high senescence score, with the small cluster 5 exhibiting high scores for both proliferation and senescence (**Figures 5D and S5B**).

Pelabresib treatment led to robust decrease in HNF1A and modest decrease in HNF4G expression (**Figure 5E-F**), consistent with in vitro data (**Figures 2A-B, 4B, 4D**). To quantify the downstream GI transcriptome, we assigned the previously defined HNF and AR scores to each single cell in both treatment conditions. Pelabresib treatment led to a significant decrease in the HNF score, suggesting an effect on the GI transcriptome (**Figure 5G**). In contrast, AR expression was not suppressed, and AR score did not decrease with pelabresib treatment (**Figures 5H-I**). This is consistent with bulk RNA-Seq data on 22Rv1 cells (**Figures 2D and S2A**). We next performed pseudobulk analysis pooling all single-cell transcriptomic data of each condition to identify differentially expressed genes between pelabresib and vehicle treatment. GSEA analysis of the pseudobulk data showed enrichment of PCa_GI, HNF1A, and HNF4G targets, as well as cell cycle-related gene sets in pelabresib downregulated genes. Senescence-related gene sets were enriched in pelabresib upregulated genes **(Figure S5D).** Collectively, these data indicate that BETi inhibits the GI transcriptome, inhibits proliferation, and induces senescence in GI transcriptome-positive prostate cancer.

### Combination efficacy of enzalutamide and pelabresib in AR-positive CRPC PDX models

Next, we asked whether BET inhibition could further synergize with AR inhibition in CRPC by targeting a parallel survival pathway of the GI transcriptome. For this we used CRPC PDXs with varied levels of HNF scores; LuCaP 70CR (AR^pos^ HNF^high^), LuCaP 77CR (AR^pos^ HNF^high^), LuCaP 35CR (AR^pos^ HNF^low^), LuCaP 145.2 (NEPC, HNF^neg^), LuCaP 49 (NEPC, HNF^neg^) and LuCaP 93 (NEPC, HNF^neg^) and treated them with enzalutamide, pelabresib and a combination of enzalutamide and pelabresib. In LuCaP 70CR, a castration resistant PDX model, enzalutamide treatment reduced tumor growth rate. Pelabresib treatment induced stronger growth inhibition and the combination of pelabresib with enzalutamide had the most potent growth inhibitory effects (**Figure 6A**). Immunoblot analysis performed on tumors collected at the end of the experiment showed a decrease in the protein level of HNF1A in tumors treated with pelabresib alone or in combination with enzalutamide. No significant change in AR protein level was detected with any of the drug treatments (**Figure 6B**). qRT-PCR analysis performed using RNA extracted from the end-of-study tumors revealed a decrease in selected GI lineage gene transcripts such as HNF1A and MUC13. Enzalutamide treatment decreased AR-target genes expression, and the combination treatment decreased both AR and HNF1A/HNF4G-target genes expression (**Figure 6C**).

**Figure 6.**
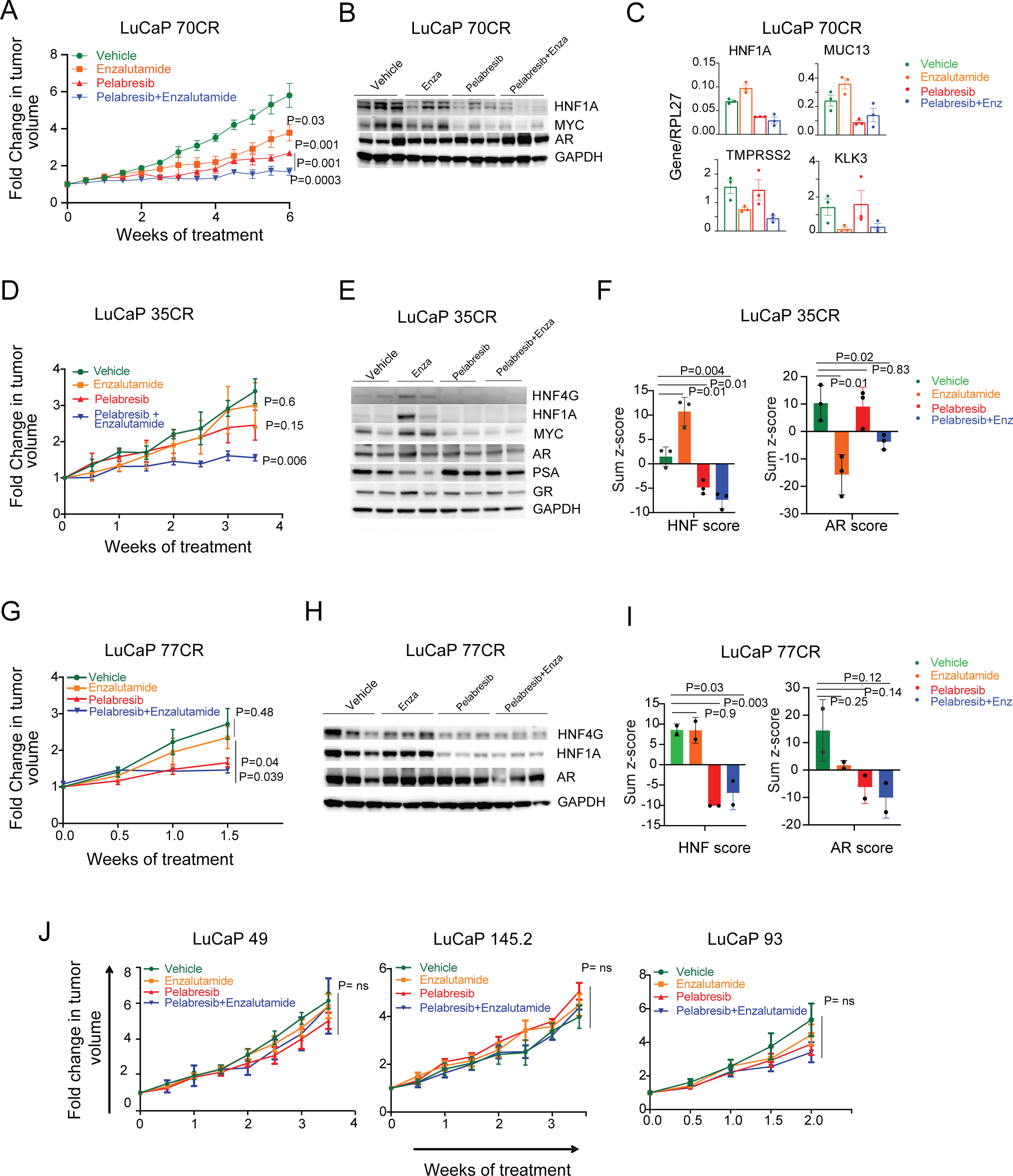
Combination efficacy of enzalutamide and pelabresib in AR-positive CRPC PDX models. (A) Treatment response of LuCaP 70CR PDX in SCID mice when treated with vehicle (0.5% methylcellulose/0.2% tween-80 in sterile water), enzalutamide (50 mg/kg), pelabresib (30 mg/kg), or enzalutamide and pelabresib. Enzalutamide and pelabresib were oral gavaged once and twice a day respectively (n=5 for all treatments). Treatment was started when tumors reached a volume of approximately 100 mm^3^. Fold change in growth rate over day 0 (start of treatment) is shown. Mean ± SEM. Two-tailed unpaired t-test. (B) Immunoblots of three representative tumor explants obtained at the end of the experiment shown in A. (C) qRT-PCR analysis of HNF1A, MUC13, TMPRSS2, and KLK3 mRNA levels in tumors harvested at the end of the study. n=3 for each treatment condition. (D) Treatment response of LuCaP 35CR PDX in SCID mice when treated with vehicle (0.5% methylcellulose/0.2% tween-80 in sterile water), enzalutamide (50 mg/kg), pelabresib (30 mg/kg), or enzalutamide and pelabresib. Enzalutamide and pelabresib were oral gavaged once and twice a day respectively (n=3 for all treatments). Treatment was started when tumors reached a volume of approximately 100 mm^3^. Fold change in growth rate over day 0 (start of treatment) is shown. Mean ± SEM. Two-tailed unpaired t-test. (E) Immunoblots of two representative tumors obtained at the end of the study shown in D. (F) Left panel shows HNF score modulation in LuCaP 35CR tumors treated with different drugs as shown in D. The HNF score was calculated using RNA-Seq gene expression generated from explanted tumors at the end of the study. The right panel shows modulation of AR signaling using the AR score. Two-tailed unpaired t-test, n=3. (G) Treatment response of LuCaP 77CR PDX in SCID mice when treated with vehicle (0.5% methylcellulose/0.2% tween-80 in sterile water), enzalutamide (50 mg/kg), pelabresib (30 mg/kg), or enzalutamide and pelabresib. Enzalutamide and pelabresib were oral gavaged once and twice a day respectively (n=3 for all treatments). Treatment was started when tumors reached a volume of approximately 100 mm^3^. Fold change in growth rate over day 0 (start of treatment) is shown. Mean ± SEM. Two-tailed unpaired t-test. (H) Immunoblots of three representative tumors obtained at the end of the study shown in G. (I) HNF score (left) and AR score (right) modulation in LuCaP 77CR tumors treated with different drugs as shown in G. Two-tailed unpaired t-test, n=2. (J) Treatment response of LuCaP 49, LuCaP 145.2, and LuCaP 93 PDXs in SCID mice when treated with vehicle, enzalutamide, pelabresib or enzalutamide and pelabresib. Treatment conditions were same as described in A, D, and G. n=3 for each treatment condition in each PDX line.

LuCaP 35CR is a castration resistant PDX model with a low HNF score. LuCaP 35CR tumors were treated with vehicle, enzalutamide, pelabresib, and the combination of pelabresib with enzalutamide for four weeks. Tumors showed resistance to enzalutamide treatment. Pelabresib as a single agent, had moderate response. However, the combination treatment of pelabresib and enzalutamide significantly reduced tumor growth (**Figure 6D**). Immunoblot analysis on protein lysates from end-of-study tumors revealed that enzalutamide treatment led to an increase in protein levels of HNF4G, HNF1A, and MUC13 (**Figure 6E**). Previously, we have noted similar observations in LNCaP/AR tumors treated with enzalutamide (Shukla et al. 2017). Importantly, the increase in GI gene expression induced by enzalutamide treatment could be effectively inhibited by combining pelabresib with enzalutamide (**Figure 6E**). RNA-Seq analysis was performed on LuCaP 35CR tumors to study global transcriptome changes under different treatment conditions. Enzalutamide treatment significantly increased the HNF score. Pelabresib decreased HNF1A expression and the HNF score, and in combination with enzalutamide, reversed the enzalutamide-induced increase in the HNF score (**Figures 6F and S6A**). The AR score decreased with enzalutamide alone and with enzalutamide and pelabresib combination treatment but not with pelabresib treatment alone (**Figures 6F and S6A**).

Similar observations were noted when we performed a short-term treatment study using LuCaP 77CR, a castration resistant, a high HNF score, and an AR-positive model. Enzalutamide alone did not cause any significant growth inhibition. In contrast, pelabresib treatment either alone or in combination with enzalutamide significantly reduced the growth of LuCaP 77CR (**Figure 6G**). Immunoblot analysis of protein lysates prepared from end-of-study tumors revealed that pelabresib treatment either alone or in combination with enzalutamide led to a decrease in protein levels of HNF4G and HNF1A. AR protein level remained unchanged under all treatment conditions (**Figure 6H**). RNA-Seq analysis performed on end-of-study tumors revealed that the HNF1A expression as well as the HNF score decreased significantly with both the pelabresib and the combination treatment while neither AR expression nor the AR score significantly altered with any of the treatments (**Figures 6I and S6B).**

Taken together, across different GI transcriptome-expressing CRPC PDX models, consistent pelabresib-mediated growth inhibition was observed. Importantly, global transcriptome analysis consistently showed robust downregulation of HNF1A, HNF4G and the HNF score with pelabresib treatment in all the PDX models assayed. Furthermore, tumor growth of the GI transcriptome expression negative PDX models (LuCaP 49, LuCaP 145.2 and LuCaP 93) was not inhibited by pelabresib treatment (**Figure 6J**). Taken together, these data suggest a selective growth inhibitory effect of BET inhibitors on GI transcriptome-expressing models.

## DISCUSSION

In prostate cancer, lineage plasticity results in extensive reprogramming of the epigenetic landscape, including changes in the cistrome of the master transcription factors FOXA1 (Baca et al. 2021) or switch to other master transcription factors, such as loss of AR and gain of ASCL1 or NEUROD1 in neuroendocrine prostate cancer (Cejas et al. 2021; Chen et al. 2023). We have uncovered that aberrant upregulation of gastrointestinal master regulators HNF4G and HNF1A alters enhancer landscape and chromatin accessibility conducive to the expression of GI-specific transcriptome in prostate cancer cells. In the present study, we found that a high GI transcriptome expression in mCRPC tumors is predictive of poor response to AR-targeted therapies and a shorter overall patients’ survival. Our previous studies have shown that genetic depletion of either HNF1A or HNF4G inhibits GI transcriptome expression. Thus, we reasoned that pharmacological targeting of either HNF1A or HNF4G would be sufficient for therapeutic studies. HNF4G is an orphan nuclear receptor with no well-characterized ligand and HNF1A is a homeobox domain containing transcription factor lacking any small molecule binding pocket. Due to expected roadblocks in identifying small molecule inhibitors regulating the activity of these transcription factors, we focused on an alternative approach of inhibiting HNF1A and HNF4G transcription.

Epigenetic therapy has been proposed to target specific lineage states in prostate cancer. Prior studies have suggested that BET, P300/CBP, LSD1, EZH2, and SWI/SNF inhibitors can disrupt AR-mediated transcriptional activity (Asangani et al. 2014; Lasko et al. 2017; Faivre et al. 2020; Welti et al. 2021; Xiao et al. 2022; Li et al. 2023). Several studies have shown BRD4 as an important cofactor required for AR transcriptional activity (Asangani et al. 2014; Faivre et al. 2020). One important caveat of these studies is that cells were hormone starved in charcoal-stripped media and the addition of DHT together with BETi led to severely impaired AR target gene expression compared to the addition of DHT alone. In these studies, and consistent with our observations, the transcription of AR itself was unaffected by BET inhibition. In our studies using 22Rv1 cells, a panel of patient-derived organoids and a panel of LuCaP PDX models, BRD4 inhibition did not consistently inhibit the AR-regulated transcriptome, though it did inhibit it in LuCaP 77CR. In a clinical trial of the BETi ZEN-3694, ∼30% of patients experienced an acute rise of PSA upon starting the drug and this rise was associated with longer progression-free survival (Aggarwal et al. 2020), a feature that distinguishes BETi from ARSIs. Although, the trial could not precisely define any biomarkers predictive of ZEN-3694 response, patients with low baseline AR signaling in tumors demonstrated longer rPFS than patients with high AR signaling (median rPFS 10.4 vs. 4.3 months). A serum PSA decline of 50% or more (PSA50) with ZEN-3694 treatment was seen in only 10% of patients and was not correlated with response to treatment (Aggarwal et al. 2020). In another clinical trial using a different BETi GS-5829, only 1 out of 31 patients showed a PSA50 decline (Aggarwal et al. 2022). These data are consistent with our results and indicate that AR transcriptome is not inhibited by BETi in patients and AR-independent mechanisms may contribute to BET inhibitor response (Aggarwal et al. 2020). Taken together, our findings not only implicate the poor prognosis of GI transcriptome expressing prostate cancer but also emphasize this subset to be vulnerable to BET inhibitors-mediated growth inhibition. Therefore, our studies have important clinical implications, and we propose that a high HNF1A/HNF4G transcriptional activity in CRPC tumors is a biomarker of an aggressive, ARSI-resistant disease that can be managed by treatment with BET inhibitors.

## AUTHOR CONTRIBUTION

Experimental design: S.S., P.C., and Y.C.; Western blots and qRT-PCRs: S.S.; ChIP-Seq, RNA-Seq, Single cell-RNA-Seq: S.S; Analysis of ChIP-Seq, RNA-Seq, and scRNA-Seq: D.L.; Cell viability assays: S.S.; Bioinformatic/Biostatistics Analysis: S.S., D.L., I.O., and Y.C. Mouse experiments design and execution: E.C., H.N., J.C., G.B., W.H.C., N.T., M.P; Resources sharing: E.C., A.G., E.C., S.A., and P.T.; Pathology supervision: A.G.; Manuscript writing: S.S., and Y.C. All authors reviewed and edited the manuscript.

## ACKNOWLEDGEMENT

Pelabresib was generously provided for in vivo experiments by Constellation Pharmaceuticals. Establishment of LuCaP PDXs was supported by the Pacific Northwest Prostate Cancer SPORE (P50CA97186), the P01 NIH grant (P01CA163227), and the Institute for Prostate Cancer Research. We would like to thank the patients who generously donated tissue that made this research possible. We would also like to thank Comparative Medicine Animal Caregivers for their assistance with the LuCaP PDX work. We acknowledge the use of the Integrated Genomics Operation Core. This work was funded by the NCI Cancer Center Grant (P30CA008748), the NCI SPORE (P50CA221745 to Y.C., P.C., A.G.), the Cancer Moonshot Drug Resistance and Sensitivity Network (DRSN) (U54CA224079 to Y.C.), R01CA193837, R01CA208100, the Prostate Cancer Foundation (Y.C.), the STARR Cancer Consortium (Y.C. and P.C.)., and the Geoffrey Beene Cancer Center (Y.C. and P.C.).

## METHODS

### Sex as a biological variable

Our study exclusively examined male mice because the disease modeled is only relevant in males.

### Cell lines, Antibodies, and Reagents

22Rv1 (ATCC CRL2505) cell lines was obtained from the American Type Culture Collection (ATCC) and maintained in RPMI supplemented with 10% Fetal bovine serum (Omega), L-glutamine (2 mM), penicillin (100 U/ml), and streptomycin (100 μg/ml). Cell lines were confirmed mycoplasma free by PCR testing.

Antibodies to the following were used for Western blotting: anti-HNF4G (Perseus Proteomics, #PP-B6502A-00; 1:1000 for Western blotting), anti-HNF1A (Santa Cruj Biotechnology, sc6547X; 1:1000 for Western blotting) anti-HNF1A (Cell Signaling Technology Cat# 89670 (also 89670S), RRID:AB_2728751; 1:1000 for Western blotting), anti-AKR1C3 (Sigma-Aldrich Cat# A6229, RRID:AB_476751; 1:1000 for Western blotting), anti-GAPDH (abmgood, G041; 1:5000 for Western blotting), anti-β-actin (Cell Signaling Technology Cat# 3700; 1:1000 for Western blotting), anti-UGT2B15 (Abcam, ab154864; 1:1000 for Western blotting), anti-MYC (Cell Signaling Technology Cat# 5605, RRID:AB_1903938; 1:2000 for Western blotting), anti-AR (Abcam Cat# ab108341, RRID:AB_10865716); 1:1000 for Western blotting), anti-GR (Cell Signaling Technology Cat# 3660, RRID:AB_11179215; 1:1000 for Western blotting), anti-PSA (Cell Signaling Technology Cat# 2475, RRID:AB_2797601; 1:1000 for Western blotting), anti-BRD4 (Bethyl laboratories, A301-985A100 for ChIP-seq).

### RNA isolation and qRT-PCR

To isolate RNA from cell lines, E.Z.N.A total RNA kit (Omega) was used. To isolate RNA from xenograft tumor explants, the tumor samples were ground in 1 ml Trizol (Invitogen) using a PowerGen homogenizer (Fisher Scientific), followed by the addition of 200 µL chloroform. The samples were then centrifuged at 10,000 g for 15 minutes. The upper phase was mixed with an equal volume of 70% ethanol, and the RNA was further purified using the E.Z.N.A total RNA kit (Omega).

For qRT-PCR, RNA was reverse transcribed using the High-Capacity cDNA Reverse Transcription Kit (ABI). Power SYBR Master Mix (ABI) was used to run PCR on a ViiA7 Real Time PCR System (Life Technologies).

### Immunoblot

Cell lysates were prepared in RIPA buffer supplemented with proteinase/phosphatase inhibitor. Proteins were resolved on NuPAGE Novex 4–12% Bis-Tris Protein Gels (Life Technologies) and transferred electrophoretically onto a PVDF 0.45 μm membrane (BioRad). Membranes were blocked for 1 hour at room temperature in blocking buffer consisting of 5% milk or 1 % BSA diluted in Tris buffer saline plus 0.1% Tween 20 (TBST) and were incubated overnight at 4 °C with the primary antibodies diluted in the same buffer. After 3 washes of 10 min in TBST, membranes were incubated with secondary antibodies diluted in blocking buffer for 1 hour at room temperature. After 3 washes of 10 minutes in TBST, Enhanced Chemiluminiscence (ECL) was performed using ECL kit (Thermo Scientific, 80196).

### Cell viability assays

Cells were plated onto 96-well plates in their respective culture medium and incubated at 37 °C in an atmosphere of 5% CO2. After overnight incubation, a serial dilution of ABBV-075 was prepared and added to the plate. The cells were further incubated for 5 days, and the CellTiter-Glo assay (Promega) was then performed according to the manufacturer’s instructions to determine cell proliferation. The luminescence signal from each well was acquired using the GLOMAX 96 microplate Luminometer (Promega), and the data were analyzed using GraphPad Prism software (RRID:SCR_002798).

### Mouse procedures

CB17 SCID male mice (Charles River) were castrated and, 2 weeks after castration, were subcutaneously implanted with tumor bits of LuCaP 35CR, 70CR, 77CR. LuCaP 49, 93 and 145.2 were implanted in intact CB17 SCID male mice. When tumors exceeded 100 mm^3^, animals were randomized to control and treatment groups (n = 3–6 per group). Treatment with enzalutamide (50 mg/kg, once a day), pelabresib (30 mg/Kg, twice daily), enazalutamide and pelabresib combination or vehicle was begun at a tumor size of 100 mm^3^. Mice were treated until the end of the experiments. Tumor volumes were monitored twice weekly. The research personnel measuring tumors were blinded to the treatment group assignment of mice.

### Gene expression analysis

RNA-seq was performed by the MSKCC Integrated Genomics Operation (IGO) core facility using poly-A capture. The libraries were sequenced on an Illumina NovaSeq 6000 platform with 100 bp paired-end reads to obtain a minimum yield of 40 million reads per sample. The sequence data were processed and mapped to the human reference genome (hg38) or mouse reference genome (mm10) using STAR (RRID:SCR_004463), version 2.3 (Dobin et al. 2013). Gene expression was quantified as transcripts per million (TPM) using “edge” R package (Robinson, McCarthy, and Smyth 2010) and log2 transformed. GSEA was performed using JAVA GSEA 2.0 program, using a difference of mean between replicates and gene permutation (Subramanian et al. 2005). The gene sets used were the Broad Molecular Signatures Database gene sets v7, c2 (curated gene sets), c5 (gene ontology gene sets), c6 (oncogenic signatures), c7 (immunologic signatures) as well as custom gene sets generated by us.

### Single Cell RNA-seq

Subcutaneous PDX tumors were harvested after vehicle or pelabresib treatment (n=2 mice for each condition). The tumors were dissociated into single-cell suspension using the tumor dissociation kit (Miltenyi Biotec, 130-095-929) following the manufacturer’s protocol. Live DAPI-negative, single tumor cells were sorted out by flow cytometry. For each sample, 5,000 cells were directly processed with 10X genomics Chromium Single Cell 3’ GEM, Library & Gel Bead Kit v3 according to the manufacturer’s specifications. For each sample, 200 million reads were acquired on NovaSeq platform S4 flow cell.

Reads obtained from the 10x Genomics scRNAseq platform were mapped to human and mouse combined genome (GRCh38 + mm10) using Cell Ranger (10X Genomics). Human and mouse cells were separated based on the ratio of mapped reads to each genome. Cells were clustered as human or mouse if they had more than 75% mapped reads with human or mouse origin. The raw sequencing fastq data of human and mouse cells were separated using the cell barcodes. The separated human and mouse fastq data were then mapped to human or mouse specific genomes separately using Cell Ranger and downstream analysis was performed separately.

True cells were distinguished from empty droplets using scCB2 package (Ni, Chen, Brown, and Kendziorski 2020). The levels of mitochondrial reads and numbers of unique molecular identifiers (UMIs) were similar among the samples, which indicates that there were no systematic biases in the libraries prepared from mice with different treatment conditions. Cells were removed if they had less than 2000 total counts, more than 60,000 total counts, less than 100 total genes, or greater than 20% mitochondrial reads. Genes detected in less than 20 cells and all mitochondrial genes were removed for subsequent analyses. Putative doublets were removed using the Doublet Detection package (Xi and Li 2021). The average gene detection in each cell type was similar among the samples. Combining human cells in the entire cohort of pelabresib and vehicle groups yielded a filtered count matrix of 17,475 cells by 23,050 genes, with a median of 12,517 counts and a median of 3,711 genes per cell. The count matrix was then normalized by counts per ten thousand (CP10K), and log(X+1) transformed for analysis of the combined dataset. The top 2000 highly variable genes were found using SCANPY (version 1.6.1) (Wolf, Angerer, and Theis 2018). Principal Component Analysis (PCA) was performed on the 2000 most variable genes with the top 50 principal components (PCs) retained with 25% variance explained. To visualize single cells of the global atlas, we used Uniform Manifold Approximation and Projection for Dimension Reduction (UMAP) (Becht et al. 2018). We then performed Leiden clustering and found 5 clusters (Traag, Waltman, and van Eck 2019). Marker genes for each cluster were found with scanpy.tl.rank_genes_groups. Cell types were determined using a combination of marker genes identified from the literature and gene ontology for cell types using the web-based tool Panglao DB (Franzen, Gan, and Bjorkegren 2019). Hierarchical clustering and heat-map generation were performed for single cells based on log-normalized and scaled expression values of marker genes curated from literature or identified as highly differentially expressed.

Differentially expressed genes between different clusters were found using MAST package (Finak et al. 2015), which were shown in heatmap. The log FC of MAST output was used for the ranked gene list in GSEA analysis (Subramanian et al. 2005).

Gene imputation was performed using MAGIC (Markov affinity-based graph imputation of cells) package, and imputed gene expression were used in theviolin plots, UMAPS, and the heatmap in Figures 5E-F, 5H, S5A and S5C (van Dijk et al. 2018).

Differentially expressed genes between pelabresib and vehicle treated samples were found using ‘FindMarkers’ function of Seurat package (Hao et al. 2021). The logFC of ‘FindMarkers’ output was used for the ranked gene list in GSEA analysis (Subramanian et al. 2005).

### Chromatin Immunoprecipitation and Sequencing

Chromatin isolation from cell lines and immunoprecipitation was performed following the protocol previously described (Shukla et al. 2017). Briefly, chromatin was isolated from 22Rv1 cells expressing HNF4G or GFP and treated with DMSO or ABBV-075. BRD4 ChIP were performed using the antibody described in the reagents section. Input DNA was also sequenced. Next-generation sequencing was performed on an Illumina NovaSeq 6000 platform with 100 bp paired-end reads. Reads were aligned to the human genome (GRCh38) using the Bowtie2 alignment software (Langmead, Trapnell, Pop, and Salzberg 2009; Langmead and Salzberg 2012). Peaks that overlapped with known blacklisted regions (Amemiya, Kundaje, and Boyle 2019) were excluded using Homer mergePeaks (Heinz et al. 2010). Duplicate reads were eliminated for subsequent analysis. Peak calling was performed using MACS 2.1 “callpeak” function comparing immunoprecipitated chromatin with input chromatin, using standard parameters and a q-value cutoff of 10^-2^ (Zhang et al. 2008). To generate a set of BRD4 peaks in all conditions, we used Homer “mergePeaks -d given” of peaks with HNF4G and GFP overexpression with DMSO and ABBV-075 treatment (Heinz et al. 2010). We subsequently merged BRD4 peaks with AR and HNF4G peaks from our prior work (GSM2277166 and GSM2277158) to generate a set of all peaks. To generate counts per peak, we used featureCounts (Liao, Smyth, and Shi 2014). We normalized the BRD4 ChIP-seq counts between HNF4G and GFP infected cells to have the same median overall peak counts. We next separated the peaks into promoter and non-promoter (i.e. enhancer) using Homer annotate.

We separately visualized 3 peak sets: 1) Non-promoter sites containing top 1,000 AR peaks that overlap with BRD4 peaks, 2) Non-promoter sites containing top 1,000 HNF4G peaks that overlap with BRD4 peaks, and 3) BRD4 enhancer peaks that do not overlap with AR or HNF4G. The ChIP-seq profiles presented were generated using Integrated Genome Viewer (IGV) software of bigWig format files, generated using the “bamCoverage” tool from deepTool2 (Ramirez et al. 2016).

### HNF signature and HNF score

HNF signature consists of *HNF1A*, *HNF4G*, and their nine strong direct downstream targets (*AKR1C3*, *ANG*, *APOH*, *CLRN3*, *GAS2*, *METTL7B*, *MUC13*, *SGK2*, and *UGT2B15*). The nine candidate genes were chosen based on the following three criteria; 1) their expression was downregulated in 22RV1 cells (HNF1A^+^, HNF4G^+^) with HNF4G and, or HNF1A knockdown. 2) their expression was upregulated in LNCaP cells (HNF1A^-^, HNF4G^-^) cells with exogenous expression of HNF4G and, or HNF1A. 3) ChIP-seq showed a direct binding of either HNF1A and, or HNF4G at their genomic loci (GSE85559 and unpublished data). An HNF score is derived from the summed z-scores of HNF signature genes expression.

### AR score

Two previously defined AR signatures (10-gene AR signature and Hieronymus AR Signature) were combined to generate a broader AR signature (Bluemn et al. 2017; Hieronymus et al. 2006). The AR score is the summed z-scores of AR signature genes expression.

### HNF4G direct targets

The direct targets of HNF4G were determined by the integration of HNF4G ChIP-Sequencing and global gene expression profiling upon HNF4G downregulation in 22Rv1 cells (GSE85559). For the analysis performed in figure 3A-B, we chose 31 genes with strong HNF4G binding at their genomic loci that also showed robust decrease in expression upon HNF4G knockdown.

### Data availability

Gene Expression Omnibus (GEO) (RRID:SCR_005012) Accession Numbers of Datasets Generated:

- GSE253805: RNA-Seq expression profile of CRPC PDX LuCaP 77CR with BET inhibitor pelabresib and AR inhibitor enzalutamide treatment.
- GSE253806.: scRNA-Seq expression profile of CRPC PDX LuCaP 70CR with BET inhibitor pelabresib treatment.
- GSE254665: RNA-Seq expression profile of CRPC PDX LuCaP 35CR when treated with BET inhibitor pelabresib and AR inhibitor enzalutamide.
- GSE254733: RNA-Seq expression profile of 22Rv1 cells with GFP or HNF4G exogenous expression when treated with BET inhibitor ABBV-075.
- GSE254869: BRD4 ChIP-Seq in 22Rv1 cells exogenously expressing HNF4G or GFP and treated with BET inhibitor ABBV-075.
- GSE254870: BET inhibitor ABBV-075 perturbed pathways in prostate cancer organoid MSK-PCa17.

## REFERENCES

Abida, W., J. Cyrta, G. Heller, D. Prandi, J. Armenia, I. Coleman, M. Cieslik, M. Benelli, D. Robinson, E. M. Van Allen, A. Sboner, T. Fedrizzi, J. M. Mosquera, B. D. Robinson, N. De Sarkar, L. P. Kunju, S. Tomlins, Y. M. Wu, D. Nava Rodrigues, M. Loda, A. Gopalan, V. E. Reuter, C. C. Pritchard, J. Mateo, D. Bianchini, S. Miranda, S. Carreira, P. Rescigno, J. Filipenko, J. Vinson, R. B. Montgomery, H. Beltran, E. I. Heath, H. I. Scher, P. W. Kantoff, M. E. Taplin, N. Schultz, J. S. deBono, F. Demichelis, P. S. Nelson, M. A. Rubin, A. M. Chinnaiyan, and C. L. Sawyers. 2019. ‘Genomic correlates of clinical outcome in advanced prostate cancer’, Proc Natl Acad Sci U S A, 116: 11428–36.

Aggarwal, R. R., M. T. Schweizer, D. M. Nanus, A. J. Pantuck, E. I. Heath, E. Campeau, S. Attwell, K. Norek, M. Snyder, L. Bauman, S. Lakhotia, F. Y. Feng, E. J. Small, W. Abida, and J. J. Alumkal. 2020. ‘A Phase Ib/IIa Study of the Pan-BET Inhibitor ZEN-3694 in Combination with Enzalutamide in Patients with Metastatic Castration-resistant Prostate Cancer’, Clinical Cancer Research, 26: 5338–47.

Aggarwal, R., A. N. Starodub, B. D. Koh, G. Xing, A. J. Armstrong, and M. A. Carducci. 2022. ‘Phase Ib Study of the BET Inhibitor GS-5829 as Monotherapy and Combined with Enzalutamide in Patients with Metastatic Castration-Resistant Prostate Cancer’, Clinical Cancer Research, 28: 3979–89.

Albrecht, B. K., V. S. Gehling, M. C. Hewitt, R. G. Vaswani, A. Cote, Y. Leblanc, C. G. Nasveschuk, S. Bellon, L. Bergeron, R. Campbell, N. Cantone, M. R. Cooper, R. T. Cummings, H. Jayaram, S. Joshi, J. A. Mertz, A. Neiss, E. Normant, M. O’Meara, E. Pardo, F. Poy, P. Sandy, J. Supko, R. J. Sims, 3rd, J. C. Harmange, A. M. Taylor, and J. E. Audia. 2016. ‘Identification of a Benzoisoxazoloazepine Inhibitor (CPI-0610) of the Bromodomain and Extra-Terminal (BET) Family as a Candidate for Human Clinical Trials’, Journal of Medicinal Chemistry, 59: 1330–9.

Alumkal, J. J., D. Sun, E. Lu, T. M. Beer, G. V. Thomas, E. Latour, R. Aggarwal, J. Cetnar, C. J. Ryan, S. Tabatabaei, S. Bailey, C. B. Turina, D. A. Quigley, X. Guan, A. Foye, J. F. Youngren, J. Urrutia, J. Huang, A. S. Weinstein, V. Friedl, M. Rettig, R. E. Reiter, D. E. Spratt, M. Gleave, C. P. Evans, J. M. Stuart, Y. Chen, F. Y. Feng, E. J. Small, O. N. Witte, and Z. Xia. 2020. ‘Transcriptional profiling identifies an androgen receptor activity-low, stemness program associated with enzalutamide resistance’, Proc Natl Acad Sci U S A, 117: 12315–23.

Amemiya, H. M., A. Kundaje, and A. P. Boyle. 2019. ‘The ENCODE Blacklist: Identification of Problematic Regions of the Genome’, Sci Rep, 9: 9354.

Asangani, I. A., V. L. Dommeti, X. Wang, R. Malik, M. Cieslik, R. Yang, J. Escara-Wilke, K. Wilder-Romans, S. Dhanireddy, C. Engelke, M. K. Iyer, X. Jing, Y. M. Wu, X. Cao, Z. S. Qin, S. Wang, F. Y. Feng, and A. M. Chinnaiyan. 2014. ‘Therapeutic targeting of BET bromodomain proteins in castration-resistant prostate cancer’, Nature, 510: 278–82.

Baca, S. C., D. Y. Takeda, J. H. Seo, J. Hwang, S. Y. Ku, R. Arafeh, T. Arnoff, S. Agarwal, C. Bell, E. O’Connor, X. Qiu, S. A. Alaiwi, R. I. Corona, M. A. S. Fonseca, C. Giambartolomei, P. Cejas, K. Lim, M. He, A. Sheahan, A. Nassar, J. E. Berchuck, L. Brown, H. M. Nguyen, I. M. Coleman, A. Kaipainen, N. De Sarkar, P. S. Nelson, C. Morrissey, K. Korthauer, M. M. Pomerantz, L. Ellis, B. Pasaniuc, K. Lawrenson, K. Kelly, A. Zoubeidi, W. C. Hahn, H. Beltran, H. W. Long, M. Brown, E. Corey, and M. L. Freedman. 2021. ‘Reprogramming of the FOXA1 cistrome in treatment-emergent neuroendocrine prostate cancer’, Nat Commun, 12: 1979.

Becht, E., L. McInnes, J. Healy, C. A. Dutertre, I. W. H. Kwok, L. G. Ng, F. Ginhoux, and E. W. Newell. 2018. ‘Dimensionality reduction for visualizing single-cell data using UMAP’, Nature Biotechnology.

Beltran, H., D. Prandi, J. M. Mosquera, M. Benelli, L. Puca, J. Cyrta, C. Marotz, E. Giannopoulou, B. V. Chakravarthi, S. Varambally, S. A. Tomlins, D. M. Nanus, S. T. Tagawa, E. M. Van Allen, O. Elemento, A. Sboner, L. A. Garraway, M. A. Rubin, and F. Demichelis. 2016. ‘Divergent clonal evolution of castration resistant neuroendocrine prostate cancer’, Nature Medicine, 22: 298–305.

Bluemn, E. G., I. M. Coleman, J. M. Lucas, R. T. Coleman, S. Hernandez-Lopez, R. Tharakan, D. Bianchi-Frias, R. F. Dumpit, A. Kaipainen, A. N. Corella, Y. C. Yang, M. D. Nyquist, E. Mostaghel, A. C. Hsieh, X. Zhang, E. Corey, L. G. Brown, H. M. Nguyen, K. Pienta, M. Ittmann, M. Schweizer, L. D. True, D. Wise, P. S. Rennie, R. L. Vessella, C. Morrissey, and P. S. Nelson. 2017. ‘Androgen Receptor Pathway-Independent Prostate Cancer Is Sustained through FGF Signaling’, Cancer Cell, 32: 474–89 e6.

Cai, L., Y. H. Tsai, P. Wang, J. Wang, D. Li, H. Fan, Y. Zhao, R. Bareja, R. Lu, E. M. Wilson, A. Sboner, Y. E. Whang, D. Zheng, J. S. Parker, H. S. Earp, and G. G. Wang. 2018. ‘ZFX Mediates Non-canonical Oncogenic Functions of the Androgen Receptor Splice Variant 7 in Castrate-Resistant Prostate Cancer’, Molecular Cell, 72: 341–54 e6.

Cejas, P., Y. Xie, A. Font-Tello, K. Lim, S. Syamala, X. Qiu, A. K. Tewari, N. Shah, H. M. Nguyen, R. A. Patel, L. Brown, I. Coleman, W. M. Hackeng, L. Brosens, K. M. A. Dreijerink, L. Ellis, S. A. Alaiwi, J. H. Seo, S. Baca, H. Beltran, F. Khani, M. Pomerantz, A. Dall’Agnese, J. Crowdis, E. M. Van Allen, J. Bellmunt, C. Morrisey, P. S. Nelson, J. DeCaprio, A. Farago, N. Dyson, B. Drapkin, X. S. Liu, M. Freedman, M. C. Haffner, E. Corey, M. Brown, and H. W. Long. 2021. ‘Subtype heterogeneity and epigenetic convergence in neuroendocrine prostate cancer’, Nat Commun, 12: 5775.

Chen, C. C., W. Tran, K. Song, T. Sugimoto, M. B. Obusan, L. Wang, K. M. Sheu, D. Cheng, L. Ta, G. Varuzhanyan, A. Huang, R. Xu, Y. Zeng, A. Borujerdpur, N. A. Bayley, M. Noguchi, Z. Mao, C. Morrissey, E. Corey, P. S. Nelson, Y. Zhao, J. Huang, J. W. Park, O. N. Witte, and T. G. Graeber. 2023. ‘Temporal evolution reveals bifurcated lineages in aggressive neuroendocrine small cell prostate cancer trans-differentiation’, Cancer Cell, 41: 2066–82 e9.

Dobin, A., C. A. Davis, F. Schlesinger, J. Drenkow, C. Zaleski, S. Jha, P. Batut, M. Chaisson, and T. R. Gingeras. 2013. ‘STAR: ultrafast universal RNA-seq aligner’, Bioinformatics, 29: 15–21.

Faivre, E. J., K. F. McDaniel, D. H. Albert, S. R. Mantena, J. P. Plotnik, D. Wilcox, L. Zhang, M. H. Bui, G. S. Sheppard, L. Wang, V. Sehgal, X. Lin, X. Huang, X. Lu, T. Uziel, P. Hessler, L. T. Lam, R. J. Bellin, G. Mehta, S. Fidanze, J. K. Pratt, D. Liu, L. A. Hasvold, C. Sun, S. C. Panchal, J. J. Nicolette, S. L. Fossey, C. H. Park, K. Longenecker, L. Bigelow, M. Torrent, S. H. Rosenberg, W. M. Kati, and Y. Shen. 2020. ‘Selective inhibition of the BD2 bromodomain of BET proteins in prostate cancer’, Nature, 578: 306–10.

Finak, G., A. McDavid, M. Yajima, J. Deng, V. Gersuk, A. K. Shalek, C. K. Slichter, H. W. Miller, M. J. McElrath, M. Prlic, P. S. Linsley, and R. Gottardo. 2015. ‘MAST: a flexible statistical framework for assessing transcriptional changes and characterizing heterogeneity in single-cell RNA sequencing data’, Genome Biol, 16: 278.

Franzen, O., L. M. Gan, and J. L. M. Bjorkegren. 2019. ‘PanglaoDB: a web server for exploration of mouse and human single-cell RNA sequencing data’, Database (Oxford), 2019.

Gao, D., I. Vela, A. Sboner, P. J. Iaquinta, W. R. Karthaus, A. Gopalan, C. Dowling, J. N. Wanjala, E. A. Undvall, V. K. Arora, J. Wongvipat, M. Kossai, S. Ramazanoglu, L. P. Barboza, W. Di, Z. Cao, Q. F. Zhang, I. Sirota, L. Ran, T. Y. MacDonald, H. Beltran, J. M. Mosquera, K. A. Touijer, P. T. Scardino, V. P. Laudone, K. R. Curtis, D. E. Rathkopf, M. J. Morris, D. C. Danila, S. F. Slovin, S. B. Solomon, J. A. Eastham, P. Chi, B. Carver, M. A. Rubin, H. I. Scher, H. Clevers, C. L. Sawyers, and Y. Chen. 2014. ‘Organoid cultures derived from patients with advanced prostate cancer’, Cell, 159: 176–87.

Han, H., Y. Wang, J. Curto, S. Gurrapu, S. Laudato, A. Rumandla, G. Chakraborty, X. Wang, H. Chen, Y. Jiang, D. Kumar, E. G. Caggiano, M. Capogiri, B. Zhang, Y. Ji, S. N. Maity, M. Hu, S. Bai, A. M. Aparicio, E. Efstathiou, C. J. Logothetis, N. Navin, N. M. Navone, Y. Chen, and F. G. Giancotti. 2022. ‘Mesenchymal and stem-like prostate cancer linked to therapy-induced lineage plasticity and metastasis’, Cell Rep, 39: 110595.

Hao, Y., S. Hao, E. Andersen-Nissen, W. M. Mauck, 3rd, S. Zheng, A. Butler, M. J. Lee, A. J. Wilk, C. Darby, M. Zager, P. Hoffman, M. Stoeckius, E. Papalexi, E. P. Mimitou, J. Jain, A. Srivastava, T. Stuart, L. M. Fleming, B. Yeung, A. J. Rogers, J. M. McElrath, C. A. Blish, R. Gottardo, P. Smibert, and R. Satija. 2021. ’Integrated analysis of multimodal single-cell data’, Cell, 184: 3573–87 e29.

Heinz, S., C. Benner, N. Spann, E. Bertolino, Y. C. Lin, P. Laslo, J. X. Cheng, C. Murre, H. Singh, and C. K. Glass. 2010. ‘Simple combinations of lineage-determining transcription factors prime cis-regulatory elements required for macrophage and B cell identities’, Molecular Cell, 38: 576–89.

Hieronymus, H., J. Lamb, K. N. Ross, X. P. Peng, C. Clement, A. Rodina, M. Nieto, J. Du, K. Stegmaier, S. M. Raj, K. N. Maloney, J. Clardy, W. C. Hahn, G. Chiosis, and T. R. Golub. 2006. ‘Gene expression signature-based chemical genomic prediction identifies a novel class of HSP90 pathway modulators’, Cancer Cell, 10: 321–30.

Labrecque, M. P., I. M. Coleman, L. G. Brown, L. D. True, L. Kollath, B. Lakely, H. M. Nguyen, Y. C. Yang, R. M. G. da Costa, A. Kaipainen, R. Coleman, C. S. Higano, E. Y. Yu, H. H. Cheng, E. A. Mostaghel, B. Montgomery, M. T. Schweizer, A. C. Hsieh, D. W. Lin, E. Corey, P. S. Nelson, and C. Morrissey. 2019. ‘Molecular profiling stratifies diverse phenotypes of treatment-refractory metastatic castration-resistant prostate cancer’, Journal of Clinical Investigation, 129: 4492–505.

Lam, H. M., R. McMullin, H. M. Nguyen, I. Coleman, M. Gormley, R. Gulati, L. G. Brown, S. K. Holt, W. Li, D. S. Ricci, K. Verstraeten, S. Thomas, E. A. Mostaghel, P. S. Nelson, R. L. Vessella, and E. Corey. 2017. ‘Characterization of an Abiraterone Ultraresponsive Phenotype in Castration-Resistant Prostate Cancer Patient-Derived Xenografts’, Clinical Cancer Research, 23: 2301–12.

Langmead, B., and S. L. Salzberg. 2012. ‘Fast gapped-read alignment with Bowtie 2’, Nature Methods, 9: 357–9.

Langmead, B., C. Trapnell, M. Pop, and S. L. Salzberg. 2009. ‘Ultrafast and memory-efficient alignment of short DNA sequences to the human genome’, Genome Biol, 10: R25.

Lasko, L. M., C. G. Jakob, R. P. Edalji, W. Qiu, D. Montgomery, E. L. Digiammarino, T. M. Hansen, R. M. Risi, R. Frey, V. Manaves, B. Shaw, M. Algire, P. Hessler, L. T. Lam, T. Uziel, E. Faivre, D. Ferguson, F. G. Buchanan, R. L. Martin, M. Torrent, G. G. Chiang, K. Karukurichi, J. W. Langston, B. T. Weinert, C. Choudhary, P. de Vries, J. H. Van Drie, D. McElligott, E. Kesicki, R. Marmorstein, C. Sun, P. A. Cole, S. H. Rosenberg, M. R. Michaelides, A. Lai, and K. D. Bromberg. 2017. ‘Discovery of a selective catalytic p300/CBP inhibitor that targets lineage-specific tumours’, Nature, 550: 128–32.

Li, M., M. Liu, W. Han, Z. Wang, D. Han, S. Patalano, J. A. Macoska, S. P. Balk, H. H. He, E. Corey, S. Gao, and C. Cai. 2023. ‘LSD1 Inhibition Disrupts Super-Enhancer-Driven Oncogenic Transcriptional Programs in Castration-Resistant Prostate Cancer’, Cancer Research, 83: 1684–98.

Liao, Y., G. K. Smyth, and W. Shi. 2014. ‘featureCounts: an efficient general purpose program for assigning sequence reads to genomic features’, Bioinformatics, 30: 923–30.

Mu, P., Z. Zhang, M. Benelli, W. R. Karthaus, E. Hoover, C. C. Chen, J. Wongvipat, S. Y. Ku, D. Gao, Z. Cao, N. Shah, E. J. Adams, W. Abida, P. A. Watson, D. Prandi, C. H. Huang, E. de Stanchina, S. W. Lowe, L. Ellis, H. Beltran, M. A. Rubin, D. W. Goodrich, F. Demichelis, and C. L. Sawyers. 2017. ‘SOX2 promotes lineage plasticity and antiandrogen resistance in TP53- and RB1-deficient prostate cancer’, Science, 355: 84–88.

Nguyen, H. M., R. L. Vessella, C. Morrissey, L. G. Brown, I. M. Coleman, C. S. Higano, E. A. Mostaghel, X. Zhang, L. D. True, H. M. Lam, M. Roudier, P. H. Lange, P. S. Nelson, and E. Corey. 2017. ‘LuCaP Prostate Cancer Patient-Derived Xenografts Reflect the Molecular Heterogeneity of Advanced Disease an--d Serve as Models for Evaluating Cancer Therapeutics’, Prostate, 77: 654–71.

Ni, Z., S. Chen, J. Brown, and C. Kendziorski. 2020. ‘CB2 improves power of cell detection in droplet-based single-cell RNA sequencing data’, Genome Biol, 21: 137.

Quintanal-Villalonga, Álvaro, Joseph M. Chan, Helena A. Yu, Dana Pe’er, Charles L. Sawyers, Triparna Sen, and Charles M. Rudin. 2020. ‘Lineage plasticity in cancer: a shared pathway of therapeutic resistance’, Nat. Rev. Clin. Oncol., 17: 360–71.

Ramirez, F., D. P. Ryan, B. Gruning, V. Bhardwaj, F. Kilpert, A. S. Richter, S. Heyne, F. Dundar, and T. Manke. 2016. ‘deepTools2: a next generation web server for deep-sequencing data analysis’, Nucleic Acids Research, 44: W160–5.

Robinson, D., E. M. Van Allen, Y. M. Wu, N. Schultz, R. J. Lonigro, J. M. Mosquera, B. Montgomery, M. E. Taplin, C. C. Pritchard, G. Attard, H. Beltran, W. Abida, R. K. Bradley, J. Vinson, X. Cao, P. Vats, L. P. Kunju, M. Hussain, F. Y. Feng, S. A. Tomlins, K. A. Cooney, D. C. Smith, C. Brennan, J. Siddiqui, R. Mehra, Y. Chen, D. E. Rathkopf, M. J. Morris, S. B. Solomon, J. C. Durack, V. E. Reuter, A. Gopalan, J. Gao, M. Loda, R. T. Lis, M. Bowden, S. P. Balk, G. Gaviola, C. Sougnez, M. Gupta, E. Y. Yu, E. A. Mostaghel, H. H. Cheng, H. Mulcahy, L. D. True, S. R. Plymate, H. Dvinge, R. Ferraldeschi, P. Flohr, S. Miranda, Z. Zafeiriou, N. Tunariu, J. Mateo, R. Perez-Lopez, F. Demichelis, B. D. Robinson, M. Schiffman, D. M. Nanus, S. T. Tagawa, A. Sigaras, K. W. Eng, O. Elemento, A. Sboner, E. I. Heath, H. I. Scher, K. J. Pienta, P. Kantoff, J. S. de Bono, M. A. Rubin, P. S. Nelson, L. A. Garraway, C. L. Sawyers, and A. M. Chinnaiyan. 2015. ‘Integrative clinical genomics of advanced prostate cancer’, Cell, 161: 1215–28.

Robinson, M. D., D. J. McCarthy, and G. K. Smyth. 2010. ‘edgeR: a Bioconductor package for differential expression analysis of digital gene expression data’, Bioinformatics, 26: 139–40.

Sequist, L. V., B. A. Waltman, D. Dias-Santagata, S. Digumarthy, A. B. Turke, P. Fidias, K. Bergethon, A. T. Shaw, S. Gettinger, A. K. Cosper, S. Akhavanfard, R. S. Heist, J. Temel, J. G. Christensen, J. C. Wain, T. J. Lynch, K. Vernovsky, E. J. Mark, M. Lanuti, A. J. Iafrate, M. Mino-Kenudson, and J. A. Engelman. 2011. ‘Genotypic and histological evolution of lung cancers acquiring resistance to EGFR inhibitors’, Sci Transl Med, 3: 75ra26.

Shukla, S., J. Cyrta, D. A. Murphy, E. G. Walczak, L. Ran, P. Agrawal, Y. Xie, Y. Chen, S. Wang, Y. Zhan, D. Li, E. W. P. Wong, A. Sboner, H. Beltran, J. M. Mosquera, J. Sher, Z. Cao, J. Wongvipat, R. P. Koche, A. Gopalan, D. Zheng, M. A. Rubin, H. I. Scher, and P. Chi. 2017. ‘Aberrant Activation of a Gastrointestinal Transcriptional Circuit in Prostate Cancer Mediates Castration Resistance’, Cancer Cell, 32: 792–806.e7.

Subramanian, A., P. Tamayo, V. K. Mootha, S. Mukherjee, B. L. Ebert, M. A. Gillette, A. Paulovich, S. L. Pomeroy, T. R. Golub, E. S. Lander, and J. P. Mesirov. 2005. ‘Gene set enrichment analysis: a knowledge-based approach for interpreting genome-wide expression profiles’, Proc Natl Acad Sci U S A, 102: 15545–50.

Tang, F., D. Xu, S. Wang, C. K. Wong, A. Martinez-Fundichely, C. J. Lee, S. Cohen, J. Park, C. E. Hill, K. Eng, R. Bareja, T. Han, E. M. Liu, A. Palladino, W. Di, D. Gao, W. Abida, S. Beg, L. Puca, M. Meneses, E. de Stanchina, M. F. Berger, A. Gopalan, L. E. Dow, J. M. Mosquera, H. Beltran, C. N. Sternberg, P. Chi, H. I. Scher, A. Sboner, Y. Chen, and E. Khurana. 2022. ‘Chromatin profiles classify castration-resistant prostate cancers suggesting therapeutic targets’, Science, 376: eabe1505.

Traag, V. A., L. Waltman, and N. J. van Eck. 2019. ‘From Louvain to Leiden: guaranteeing well-connected communities’, Sci Rep, 9: 5233.

van Dijk, D., R. Sharma, J. Nainys, K. Yim, P. Kathail, A. J. Carr, C. Burdziak, K. R. Moon, C. L. Chaffer, D. Pattabiraman, B. Bierie, L. Mazutis, G. Wolf, S. Krishnaswamy, and D. Pe’er. 2018. ‘Recovering Gene Interactions from Single-Cell Data Using Data Diffusion’, Cell, 174: 716–29 e27.

Watson, P. A., V. K. Arora, and C. L. Sawyers. 2015. ‘Emerging mechanisms of resistance to androgen receptor inhibitors in prostate cancer’, Nature Reviews: Cancer, 15: 701–11.

Welti, J., A. Sharp, N. Brooks, W. Yuan, C. McNair, S. N. Chand, A. Pal, I. Figueiredo, R. Riisnaes, B. Gurel, J. Rekowski, D. Bogdan, W. West, B. Young, M. Raja, A. Prosser, J. Lane, S. Thomson, J. Worthington, S. Onions, J. Shannon, S. Paoletta, R. Brown, D. Smyth, G. W. Harbottle, V. S. Gil, S. Miranda, M. Crespo, A. Ferreira, R. Pereira, N. Tunariu, S. Carreira, A. J. Neeb, J. Ning, A. Swain, D. Taddei, Su C. Pcf International Prostate Cancer Dream Team, M. J. Schiewer, K. E. Knudsen, N. Pegg, and J. S. de Bono. 2021. ‘Targeting the p300/CBP Axis in Lethal Prostate Cancer’, Cancer Discov, 11: 1118–37.

Wolf, F. A., P. Angerer, and F. J. Theis. 2018. ‘SCANPY: large-scale single-cell gene expression data analysis’, Genome Biol, 19: 15.

Xi, N. M., and J. J. Li. 2021. ‘Protocol for executing and benchmarking eight computational doublet-detection methods in single-cell RNA sequencing data analysis’, STAR Protoc, 2: 100699.

Xiao, L., A. Parolia, Y. Qiao, P. Bawa, S. Eyunni, R. Mannan, S. E. Carson, Y. Chang, X. Wang, Y. Zhang, J. N. Vo, S. Kregel, S. A. Simko, A. D. Delekta, M. Jaber, H. Zheng, I. J. Apel, L. McMurry, F. Su, R. Wang, S. Zelenka-Wang, S. Sasmal, L. Khare, S. Mukherjee, C. Abbineni, K. Aithal, M. S. Bhakta, J. Ghurye, X. Cao, N. M. Navone, A. I. Nesvizhskii, R. Mehra, U. Vaishampayan, M. Blanchette, Y. Wang, S. Samajdar, M. Ramachandra, and A. M. Chinnaiyan. 2022. ‘Targeting SWI/SNF ATPases in enhancer-addicted prostate cancer’, Nature, 601: 434–39.

Zhang, Y., T. Liu, C. A. Meyer, J. Eeckhoute, D. S. Johnson, B. E. Bernstein, C. Nusbaum, R. M. Myers, M. Brown, W. Li, and X. S. Liu. 2008. ‘Model-based analysis of ChIP-Seq (MACS)’, Genome Biol, 9: R137.

