## Supplemental Figures S1-S6 for "BET inhibitors as a therapeutic intervention in gastrointestinal gene signature-positive castration-resistant prostate cancer"

### Figure S1

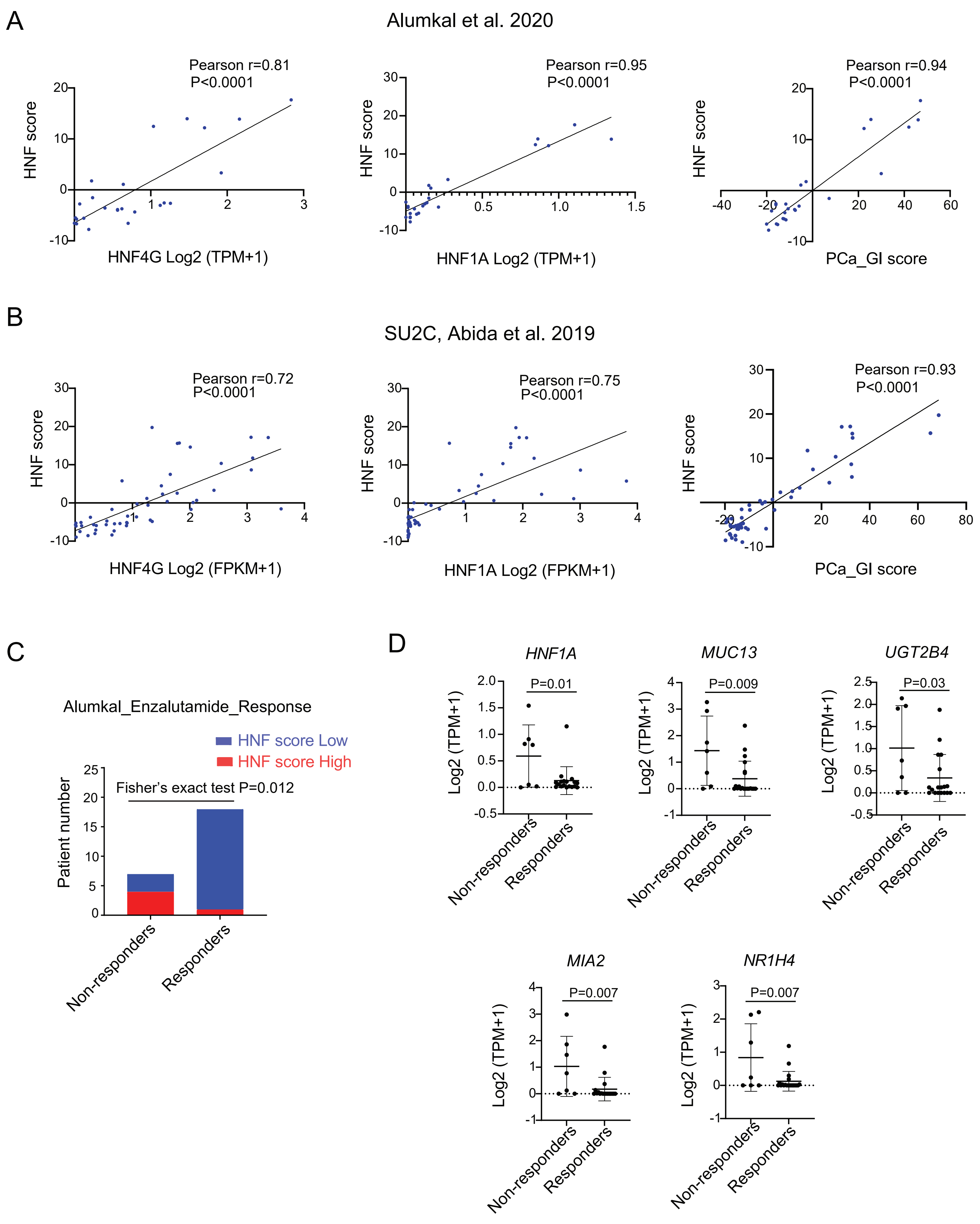
A
Alumkal et al. 2020
Pearson r=0.81
P<0.0001
HNF score
HNF4G Log2 (TPM+1)
Pearson r=0.95
P<0.0001
HNF score
HNF1A Log2 (TPM+1)
Pearson r=0.94
P<0.0001
HNF score
PCa_GI score
B
SU2C, Abida et al. 2019
Pearson r=0.72
P<0.0001
HNF score
HNF4G Log2 (FPKM+1)
Pearson r=0.75
P<0.0001
HNF score
HNF1A Log2 (FPKM+1)
Pearson r=0.93
P<0.0001
HNF score
PCa_GI score
C
Alumkal_Enzalutamide_Response
HNF score Low
HNF score High
Fisher's exact test P=0.012
Patient number
Non-responders
Responders
D
HNF1A
P=0.01
MUC13
P=0.009
UGT2B4
P=0.03
MIA2
P=0.007
NR1H4
P=0.007
Log2 (TPM+1)
Non-responders
Responders

**Figure S1. HNF score correlates with HNF1A and HNF4G expression.**

(A) Correlation of 11-gene HNF score with HNF4G transcript, HNF1A transcript, and the broader PCa_GI signature sum (Z-score) across all tumors (n=25) for which RNAseq data was available (Alumkal et al. 2020). Pearson's correlation coefficient and p value are indicated on each plot (see Figure 1). (B) Correlation of 11-gene HNF score with HNF4G transcript, HNF1A transcript and the broader PCa_GI signature sum (Z-score) across all Taxane and ARSi naive tumors (n=50) for which RNAseq data was available in the SU2C dataset (Robinson et al. 2015; Abida et al. 2019). Pearson's correlation coefficient and p value are indicated on each plot (see Figure 1). (C) HNF scores of enzalutamide non-responding and responding patient tumors in Alumkal dataset. Statistical significance is determined using Fisher’s exact test. (D) Scatter plots of selected GI-transcriptome genes expression in enzalutamide non-responding and responding tumors. Mean ± SD. Two-tailed unpaired t-test.

Figure S2

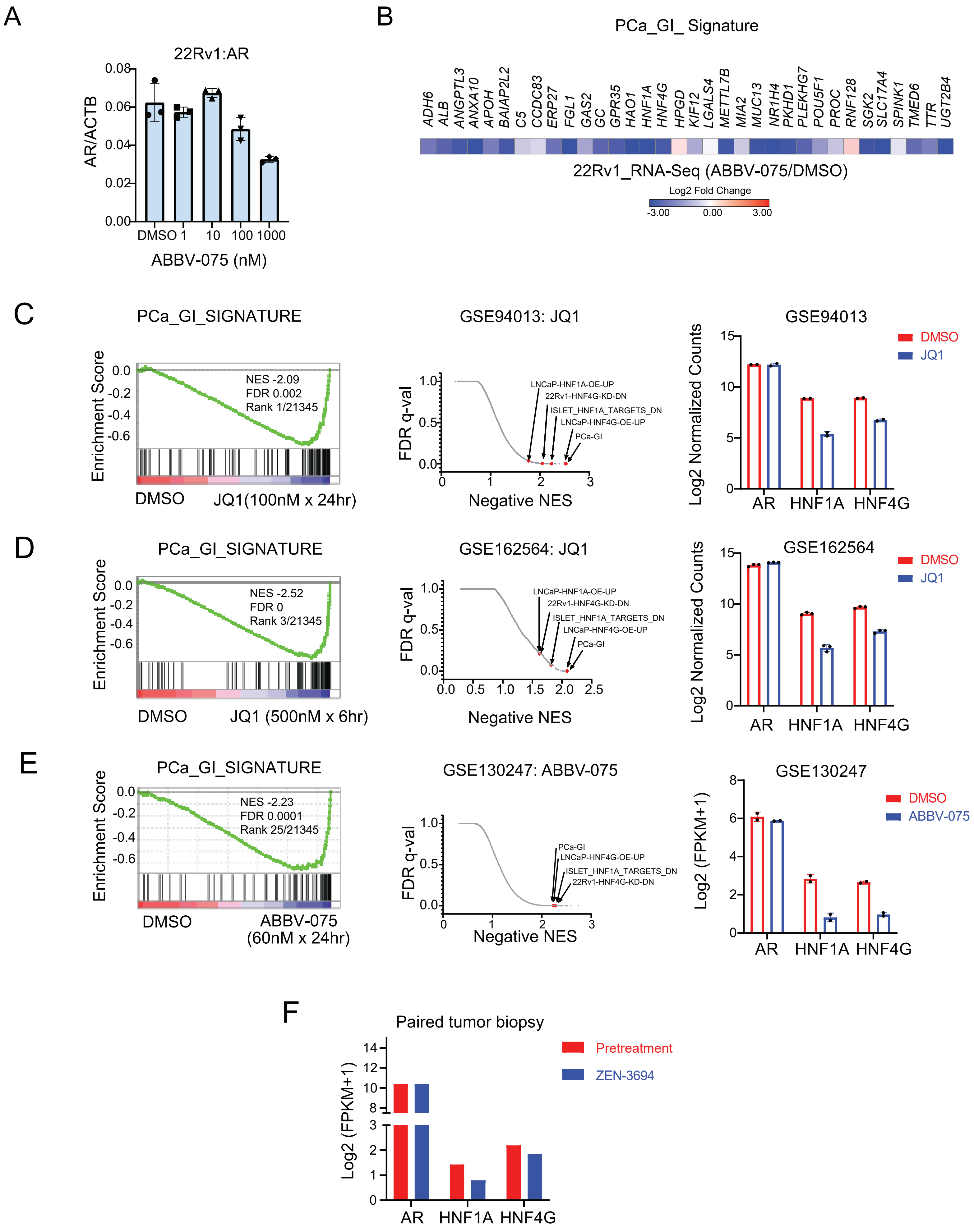
A
22Rv1:AR
AR/ACTB
0.08
0.06
0.04
0.02
0.00
DMSO 1 10 100 1000
ABBV-075 (nM)
B
PCa_GI_ Signature
ADH6 ALB ANGPTL3 ANXA10 APOH BAIAP2L2 C5 CCDC83 ERP27 FGL1 GAS2 GC GPR35 HAO1 HNF1A HNF4G HPGD KIF12 LGALS4 METTL7B MIA2 MUC13 NR1H4 PKHD1 PLEKHG7 POU5F1 PROC RNF128 SGK2 SLC17A4 SPINK1 TMED6 TTR UGT2B4
22Rv1_RNA-Seq (ABBV-075/DMSO)
Log2 Fold Change
-3.00 0.00 3.00
C
PCa_GI_SIGNATURE
Enrichment Score
NES -2.09
FDR 0.002
Rank 1/21345
DMSO JQ1(100nM x 24hr)
GSE94013: JQ1
FDR q-val
LNCaP-HNF1A-OE-UP
22Rv1-HNF4G-KD-DN
ISLET_HNF1A_TARGETS_DN
LNCaP-HNF4G-OE-UP
PCa-GI
Negative NES
GSE94013
Log2 Normalized Counts
DMSO
JQ1
AR HNF1A HNF4G
D
PCa_GI_SIGNATURE
Enrichment Score
NES -2.52
FDR 0
Rank 3/21345
DMSO JQ1 (500nM x 6hr)
GSE162564: JQ1
FDR q-val
LNCaP-HNF1A-OE-UP
22Rv1-HNF4G-KD-DN
ISLET_HNF1A_TARGETS_DN
LNCaP-HNF4G-OE-UP
PCa-GI
Negative NES
GSE162564
Log2 Normalized Counts
DMSO
JQ1
AR HNF1A HNF4G
E
PCa_GI_SIGNATURE
Enrichment Score
NES -2.23
FDR 0.0001
Rank 25/21345
DMSO ABBV-075 (60nM x 24hr)
GSE130247: ABBV-075
FDR q-val
PCa-GI
LNCaP-HNF4G-OE-UP
ISLET_HNF1A_TARGETS_DN
22Rv1-HNF4G-KD-DN
Negative NES
GSE130247
Log2 (FPKM+1)
DMSO
ABBV-075
AR HNF1A HNF4G
F
Paired tumor biopsy
Log2 (FPKM+1)
Pretreatment
ZEN-3694
AR HNF1A HNF4G

**Figure S2. BET inhibitors downregulate the GI transcriptome.**

(A) qRT-PCR of AR expression following 4 hours of treatment with ABBV-075 at indicated doses in 22Rv1 cells. (B) Heatmap of RNA-seq expression of PCa_GI signature genes in 22Rv1 cells after treatment with 25 nM ABBV-075 for 24 hours. Data is plotted as the log2 difference in gene expression between ABBV-075 and DMSO treated cells. (C-E) GSEA analysis of publicly available RNA-Seq gene expression data sets of 22RV1 cells treated with BET inhibitors JQ1 and ABBV-075. GSEA plots of PCa-GI gene signature is shown in the left panels. Middle panels show the global representation of GSEA for each experiment. X-axis shows the negative normalized enrichment score, and y-axis is the FDR q-value. HNF4G and HNF1A regulated gene sets and the PCa_GI signature gene sets are indicated by arrows. NES: Normalized enrichment score. FDR: False discovery rate. The right panels show expression of AR, HNF1A and HNF4G in each RNA-Seq experiment. (F) AR, HNF1A, and HNF4G expression in pre- and post-ZEN-3694 treated patient biopsies.

### Figure S3

A

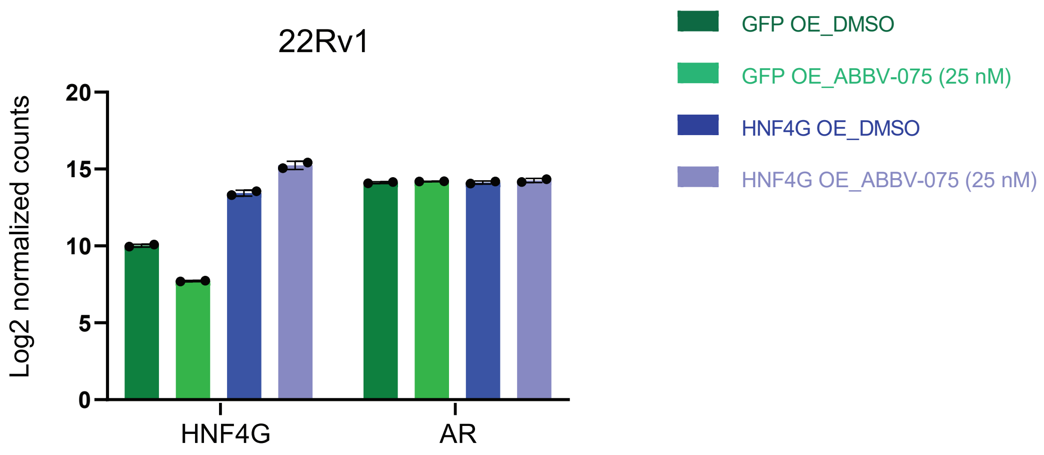


B

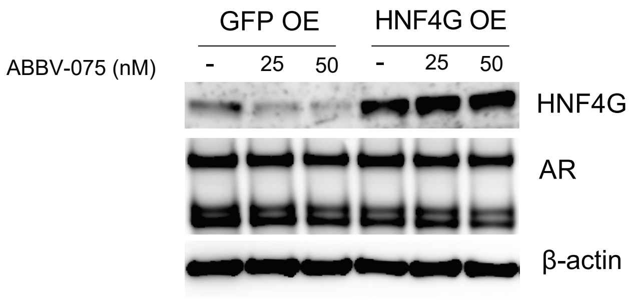


**<u>Figure S3. Expression of HNF4G from MSCV promoter is not regulated by BET proteins</u>.**

(A) Log2 normalized values of HNF4G and AR expression in 22Rv1 cells overexpressing either GFP or HNF4G from MSCV promoter when treated with DMSO or ABBV-075. (B) Immunoblot depicting HNF4G and AR protein levels in GFP and HNF4G overexpressing 22Rv1 cells treated with ABBV-075 or DMSO for 24 hours.

Figure S4

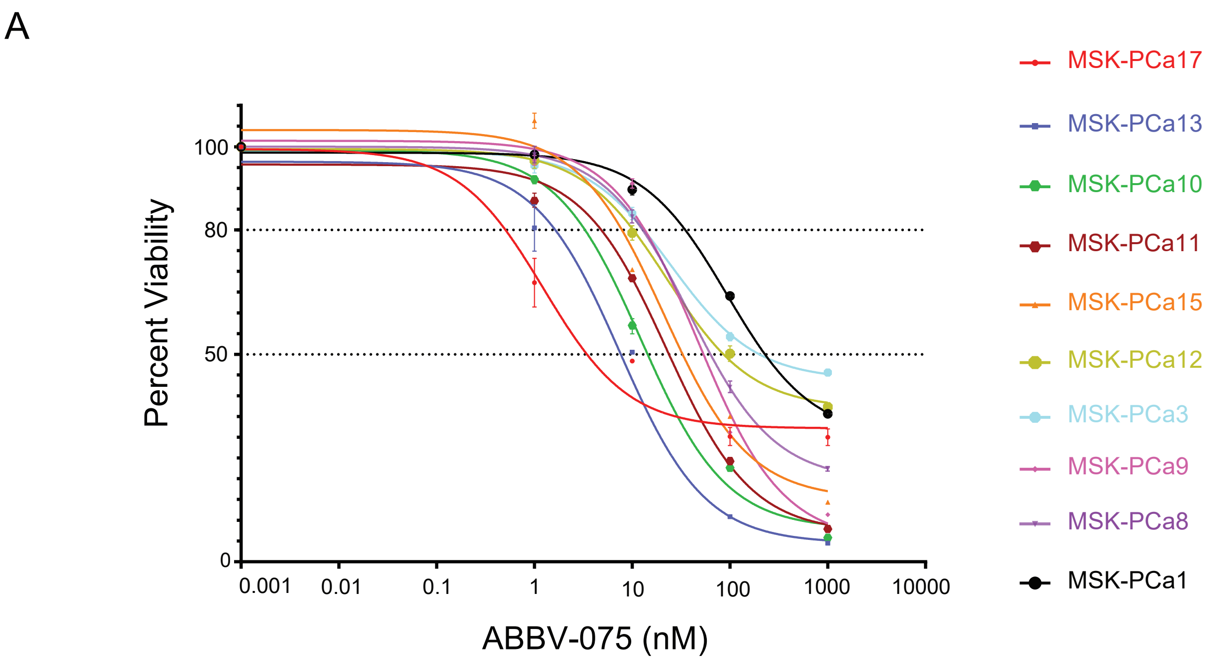


**Figure S4. IC50 curves of MSK-PCa organoids to ABBV-075 treatment.**

(A) IC50 curves showing the response of patient-derived organoids to ABBV-075 treatment. Mean ± SD (n=3).

### Figure S5

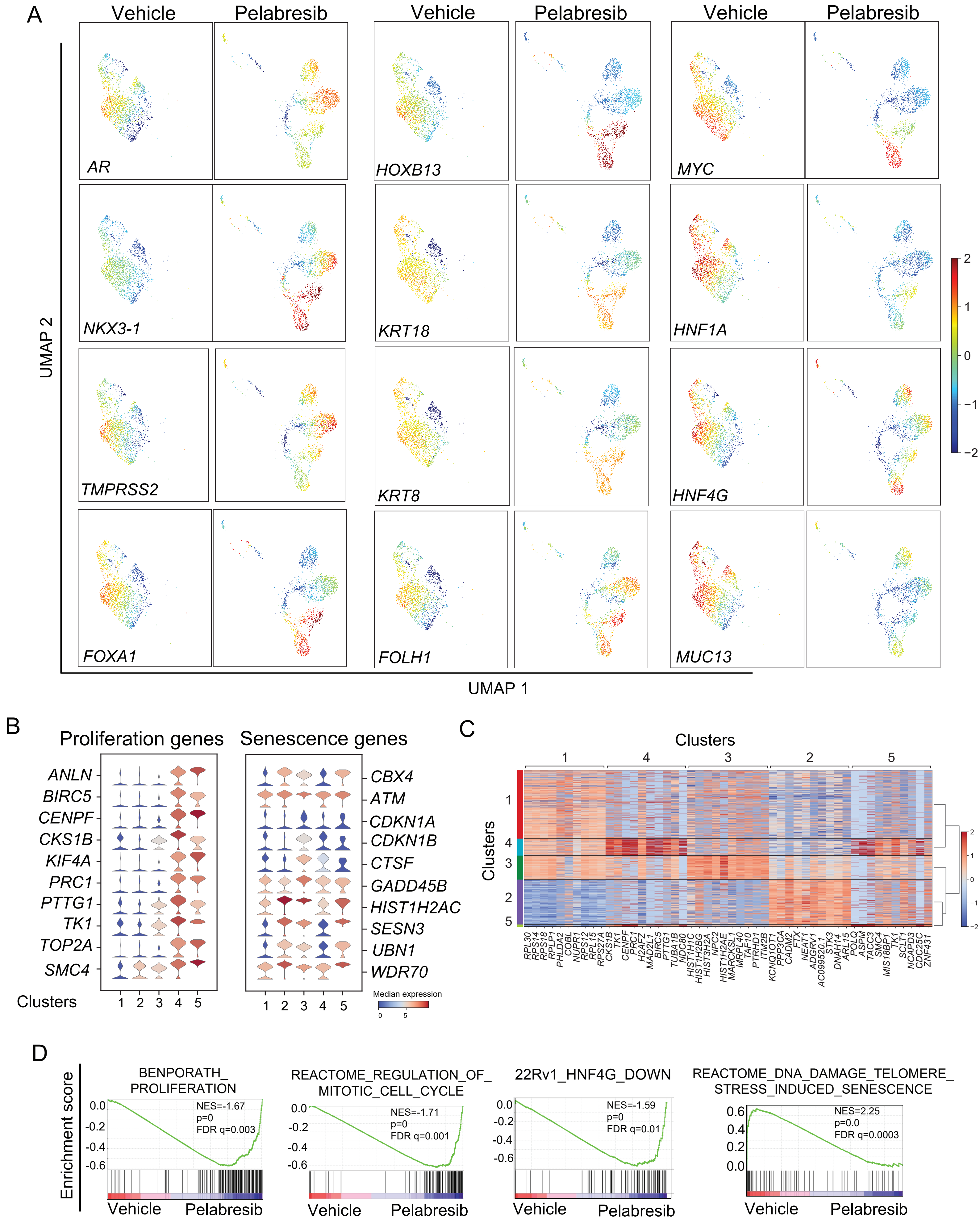

A
Vehicle
Pelabresib
Vehicle
Pelabresib
Vehicle
Pelabresib
AR
HOXB13
MYC
NKX3-1
KRT18
HNF1A
TMPRSS2
KRT8
HNF4G
FOXA1
FOLH1
MUC13
UMAP 2
UMAP 1
2
1
0
−1
−2
B
Proliferation genes
Senescence genes
ANLN
BIRC5
CENPF
CKS1B
KIF4A
PRC1
PTTG1
TK1
TOP2A
SMC4
CBX4
ATM
CDKN1A
CDKN1B
CTSF
GADD45B
HIST1H2AC
SESN3
UBN1
WDR70
Clusters
1 2 3 4 5
1 2 3 4 5
Median expression
0
5
C
Clusters
1 4 3 2 5
Clusters
1
4
3
2
5
RPL30 RPS14 RPS18 RPLP1 PHLDA2 COBL NUPR1 RPS12 RPL15 RPS27A CKS1B TK1 CENPF PRC1 H2AFZ MAD2L1 BIRC5 PTTG1 TUBA1B NDC80 HIST1H1C HIST1H2BG HIST3H2A NPC2 HIST1H2AE MARCKSL1 MRPL40 TAF10 PTRHD1 ITM2B KCNQ1OT1 PPP3CA CADM2 FTX NEAT1 ADGRV1 AC099520.1 STK3 DNAH14 ARL15 POLQ ASPM TACC3 SMC4 MIS18BP1 TK1 SCLT1 NCAPD3 CDC25C ZNF431
2
1
0
−1
−2
D
Enrichment score
BENPORATH_
PROLIFERATION
NES=-1.67
p=0
FDR q=0.003
0.0
-0.2
-0.4
-0.6
Vehicle
Pelabresib
REACTOME_REGULATION_OF_
MITOTIC_CELL_CYCLE
NES=-1.71
p=0
FDR q=0.001
0.0
-0.2
-0.4
-0.6
Vehicle
Pelabresib
22Rv1_HNF4G_DOWN
NES=-1.59
p=0
FDR q=0.01
0.0
-0.2
-0.4
-0.6
Vehicle
Pelabresib
REACTOME_DNA_DAMAGE_TELOMERE_
STRESS_INDUCED_SENESCENCE
NES=2.25
p=0.0
FDR q=0.0003
0.6
0.4
0.2
0.0
Vehicle
Pelabresib

**<u>Figure S5. Single-cell RNA-seq (scRNA-Seq) of LuCaP 70CR depicting phenotypic and transcriptomic changes with pelabresib treatment.</u>**

(A) Uniform manifold approximation and projection (UMAP) of scRNA-Seq profiles of selected genes expression in vehicle and pelabresib-treated LuCaP 70CR tumors. Color bar: z-score of log2 counts per ten thousand (log2 [CP10K+1]) (B) Violin plots of proliferation and senescence-related marker genes expression across the five clusters observed in UMAP plot of LuCaP 70CR tumors with vehicle and pelabresib treatment. Clusters 1 and 4 are enriched in vehicle-treated tumors while clusters 2, 3, and 5 are enriched with pelabresib treatment. Color in the violin plots indicates the median normalized expression level of genes in each cluster. (C) Heatmap depicting the highly differentially expressed genes (DEGs) for each cluster compared to the rest. Color bar: z-score of log2 counts per ten thousand (log2 [CP10K+1]). (D) GSEA performed on a ranked list of genes obtained from pseudo-bulk RNA-seq analysis of pooled scRNA-Seq transcriptomics data showing downregulation of proliferative and HNF4G regulated gene sets and enrichment of senescence-related gene sets in pelabresib treated cells. Please see the methods section for details.

### Figure S6

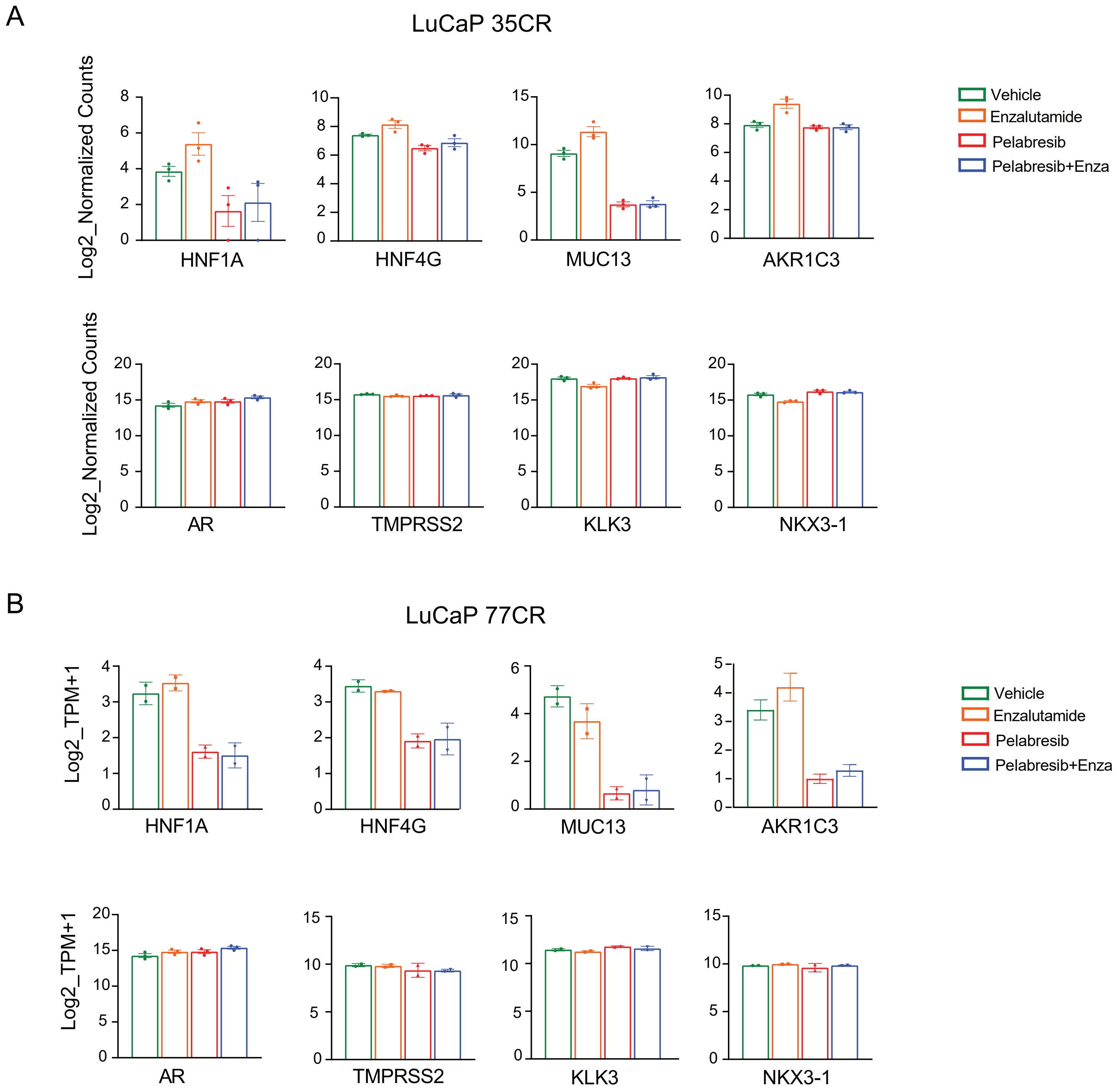


**Figure S6. Pelabresib treatment downregulates GI genes expression in LuCaP tumors.**

(A) RNA-Seq of LuCaP 35CR tumors performed with different treatment conditions showing the expression of selected GI and AR target genes. (B) RNA-Seq of LuCaP 77CR tumors performed with different treatment conditions showing the expression of selected GI and AR target genes.
